# Patterned wireless transcranial optogenetics generates artificial perception

**DOI:** 10.1101/2024.09.20.613966

**Authors:** Mingzheng Wu, Yiyuan Yang, Jinglan Zhang, Andrew I. Efimov, Abraham Vázquez-Guardado, Xiuyuan Li, Kaiqing Zhang, Yue Wang, Jianyu Gu, Liangsong Zeng, Jiaqi Liu, Mohammad Riahi, Hyoseo Yoon, Minsung Kim, Haohui Zhang, Minkyu Lee, Jiheon Kang, Kaila Ting, Stephen Cheng, Wenming Zhang, Anthony Banks, Cameron H. Good, Julia M. Cox, Lucas Pinto, Yonggang Huang, Yevgenia Kozorovitskiy, John A. Rogers

**Author notes:** Corresponding authors. (YY); (YH); (YK); (JAR). These authors contributed equally to this work (Mingzheng Wu, Yiyuan Yang, Jinglan Zhang, Andrew I. Efimov, Abraham Vázquez-Guardado, Xiuyuan Li, Kaiqing Zhang).

## Abstract

Synthesizing perceivable artificial neural inputs independent of typical sensory channels remains a fundamental challenge in the development of next-generation brain-machine interfaces. Establishing a minimally invasive, wirelessly effective, and miniaturized platform with long-term stability is crucial for creating a clinically meaningful interface capable of mediating artificial perceptual feedback. In this study, we demonstrate a miniaturized fully implantable wireless transcranial optogenetic encoder designed to generate artificial perceptions through digitized optogenetic manipulation of large cortical ensembles. This platform enables the spatiotemporal orchestration of large-scale cortical activity for remote perception genesis via real-time wireless communication and control, with optimized device performance achieved by simulation-guided methods addressing light and heat propagation during operation. Cue discrimination during operant learning demonstrates the wireless genesis of artificial percepts sensed by mice, where spatial distance across large cortical networks and sequential order-based analyses of discrimination performance reveal principles that adhere to general perceptual rules. These conceptual and technical advancements expand our understanding of artificial neural syntax and its perception by the brain, guiding the evolution of next-generation brain-machine communication.

## Main text

Creating artificial connections between the brain and the external world is becoming possible with recent advances in neuroscience and neurotechnology^1–4^. The establishment of independent input channels bypassing the typical sensory pathways enables effective brain-machine communication, allowing healthy or sensory-impaired individuals to remotely perceive senses in the extended reality^5–9^. Fundamental questions persist regarding the transmission of encoded information without external input and whether the artificial neural syntax can be meaningfully perceived by the brain. To explore these questions, reliable bio-integrated neurotechnologies incorporating wireless communication in minimally invasive implantable form factors are required to serve as functional interfaces that deliver information to the brain in real time. Current and envisioned brain-machine interfaces (BMI) consist of intricate setups that disrupt natural behavior, lack cell-type-specific capabilities, and struggle to achieve sustained long-term operation in biological systems^10,11^.

In this study, we have developed a fully implantable wireless transcranial optogenetic platform to generate artificial perceptions through spatiotemporally orchestrated optogenetic activation among distributed cortical regions in mice. This platform stands in contrast to prior elegant studies that elaborated on the processing of artificial inputs within a cortical or subcortical brain region at cellular resolution^12–14^. Higher-order cognitive behaviors are thought to result from brain-wide dynamics^15–18^, yet how distributed synthetic cortical activity is perceived remains unknown. Here, the generation of synthetic perceptions relies on large-scale sequential cortical activation, where the spatiotemporal patterns of cortical activity represent cues for mice to make decisions in an operant learning paradigm. The spatial distance among activated cortical area sequences defines the discernibility of synthesized perceptions, contributing to the perceived similarities among cortical activity patterns, captured in behavioral outcomes. This work introduces a unique wireless transcranial encoder and provides a foundational framework for expanding brain-machine communication into a broader cognitive domain through patterned cortical activations. This technology, compatible with standardized manufacturing processes in the flexible electronics industry, will permit rapid and broad dissemination to the neuroscience and BMI communities.

## Engineering platform for dynamic wireless transcranial perception modulation

The fully implantable transcranial optogenetic encoder features a wirelessly powered array of micro-scale inorganic light-emitting diodes (µ-ILEDs, 300 × 300 × 90 μm^3^) with independent, real-time wireless control over patterned optogenetic modulation of neuronal activity. The platform employs multilayer copper- polymer flexible printed circuit board (fPCB) to define the device layout and serve as the substrate for electronic components. This scheme eliminates the need for complex processes in microfabrication and it provides an immediate path to scale-up manufacture and broad distribution^19,20^ (**Extended Data Fig. 1**). As optimized for transcranial modulation of cortical activity in mice, the engineered design adopts a modular concept: a flexible optogenetic display (FOD) based on a programmable array of µ-ILEDs and a wireless control module physically separated but electrically connected through a set of mechanically compliant thin serpentine traces (**Fig. 1a**). A conformal coating of parylene-C (14 μm) symmetrically encapsulates and isolates the electronic components, FOD, and serpentine traces from the in vivo biofluidic environment. An additional coating of a soft silicone elastomer (400 μm, Young’s modulus: ∼60 kPa) creates a compliant device-tissue interface. This coating also provides further mechanical support to the serpentine connections to improve their stretchability and bendability, thereby ensuring stable operation during repetitive cycles of deformations induced by natural motions of the animal.

**Fig. 1.**
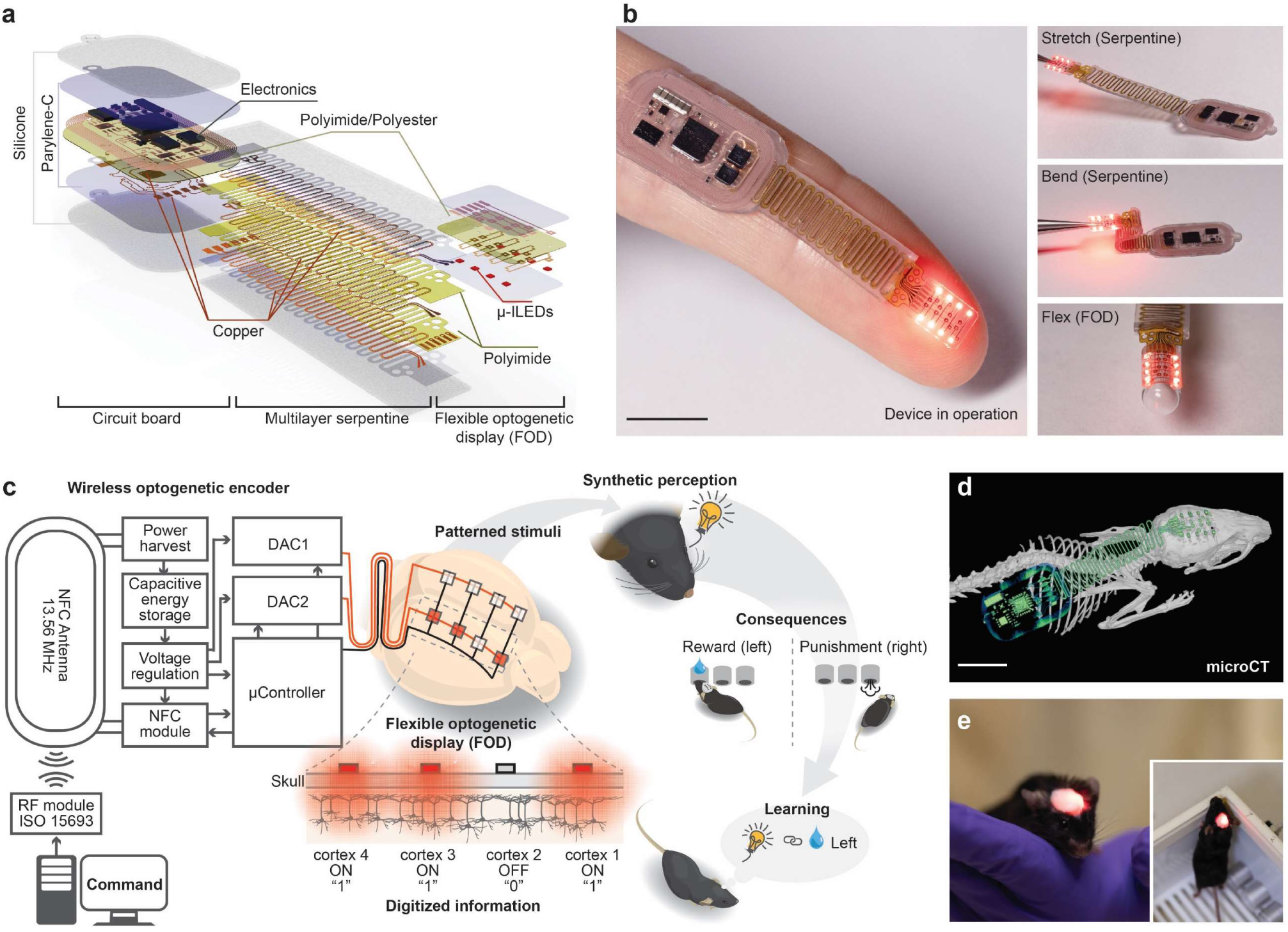
Engineering platform for dynamic wireless transcranial perception modulation. (a) Layered schematic illustration of a wireless optogenetic encoder for dynamic transcranial control of large-scale cortical activation. (b) Left, image of the flexible optogenetic display (FOD) in operation. Right, images showing the controlled patterns of µ-ILEDs during applied deformation of serpentine interconnects and FOD. Scale bar: 10 mm. (c) Schematic illustrations of the electronic modules and FOD that enable dynamic perception modulation in the context of learning. (d) Microscale computed tomographic (microCT) image of the wireless optogenetic encoder implanted in a C57/BL6J mouse. Scale bar: 10 mm. (e) A close-up photograph showing a device-integrated mouse close to an electromagnetic field resonating at 13.56 MHz. Inset, a device-integrated mouse behaving in the experimental arena circulated by a transmission antenna.

As mounted over the skull, the FOD targets selected, spatially separated groups of neurons in the cerebral cortex through transcranial optogenetic stimulation. The electronic module resides at the lumbar spine level of the animal, utilizing the broad subdermal space to house the receiver antenna and electronic components for continuous wireless power supply and dynamic independent control over individual μ-ILEDs in the FOD (**Fig. 1b**). The wireless optogenetic encoder utilizes magnetic inductive power transfer between resonant transmission and receiver antennas operating at the industrial, scientific, and medical radio frequency (ISMRF, 13.56 MHz) band, located around the experimental enclosure and device perimeter, respectively. The combination of a near-field communication (NFC) system-on-a-chip and a microcontroller, equipped with a dedicated firmware, facilitates remote ON-OFF (**Supplementary Video 1**) and intensity control over individual μ-ILEDs (**Supplementary Video 2**) using a hybridized analog/digital matrix multiplexing approach (**Fig. 1c**) through a library application programming interface in MATLAB (The MathWorks, Inc.). The encoded spatiotemporal patterns of illumination provide the brain with digitized information and programmable intensity level, adding an additional layer of freedom to control the activation volume of cortical areas. Similar to a previously reported design in a different context^21^, a collection of ceramic capacitors (5 × 22 µF, ∼1.37 mJ at 5 V) with low internal resistances enables fast discharge dynamics to provide short transcranial optical pulses (e.g., 2 ms) at high-intensity (∼70 mW/mm^2^) with rapid charge replenishment. Integration of this technology with open-source behavioral systems (Bpod, Sanworks, LLC) forms the basis of a generalizable platform for dynamically reconfigurable stimulation patterns driven by behavioral outcomes in real-time. Generation of digitized information in this manner to induce perceptual modulation can be further evaluated through diverse cue-based operant learning paradigms where mice detect or distinguish among artificial perceptions to seek external rewards (**Fig. 1c**).

The thin, subdermal implantable design enables its application in small animal models (e.g., mice, **Fig. 1d**) as a fully implanted device, without affecting their natural behaviors in complex experimental environments (**Fig. 1e**). Furthermore, iterative simulation-guided optimization of the geometries and the multilayer configurations of the serpentine interconnects ensures robust operation, without fracture or fatigue in the constituent materials (**Supplementary Methods and Extended Data Fig. 2a-i)**. The optimization increases the effective serpentine stretchability from 7% (basic design) to 28% (optimized design, **Extended Data Fig. 3a-c**). Benchtop cyclic stretching and bending tests validate these enhancements, with the conductance stability improving from 1,000 cycles (basic design) to at least 20,000 cycles (optimized design, **Extended Data Fig. 3d-f**). The longevity of the resulting devices meets practical requirements of long-term experimental protocols (**Extended Data Fig. 3g**). The overall design allows straightforward adaptations to accommodate other experimental models ranging from rodents to non-human primates. Further details on device fabrication, electronic modules, and implantation procedures appear in **Extended Data Fig. 4, Methods, and Supplementary Methods.**

## Spatiotemporal characteristics of light propagation and heat generation

A comprehensive understanding of transcranial light propagation through the brain is essential for assessing the biological implications of optical neuromodulation^22–24^. Moreover, the potential off-target opsin-independent effects, such as heat accumulation, are significant considerations in preventing unintended changes in neuronal activity^25^. Parasitic heating of the µ-ILEDs and light absorption contribute to thermal stress in the adjacent and illuminated regions (**Fig. 2a**). Operation of multiple µ-ILEDs leads to additive effects in the spatiotemporal domain, requiring additional consideration in the geometric design of the devices and stimulation parameters in experiments.

**Fig. 2.**
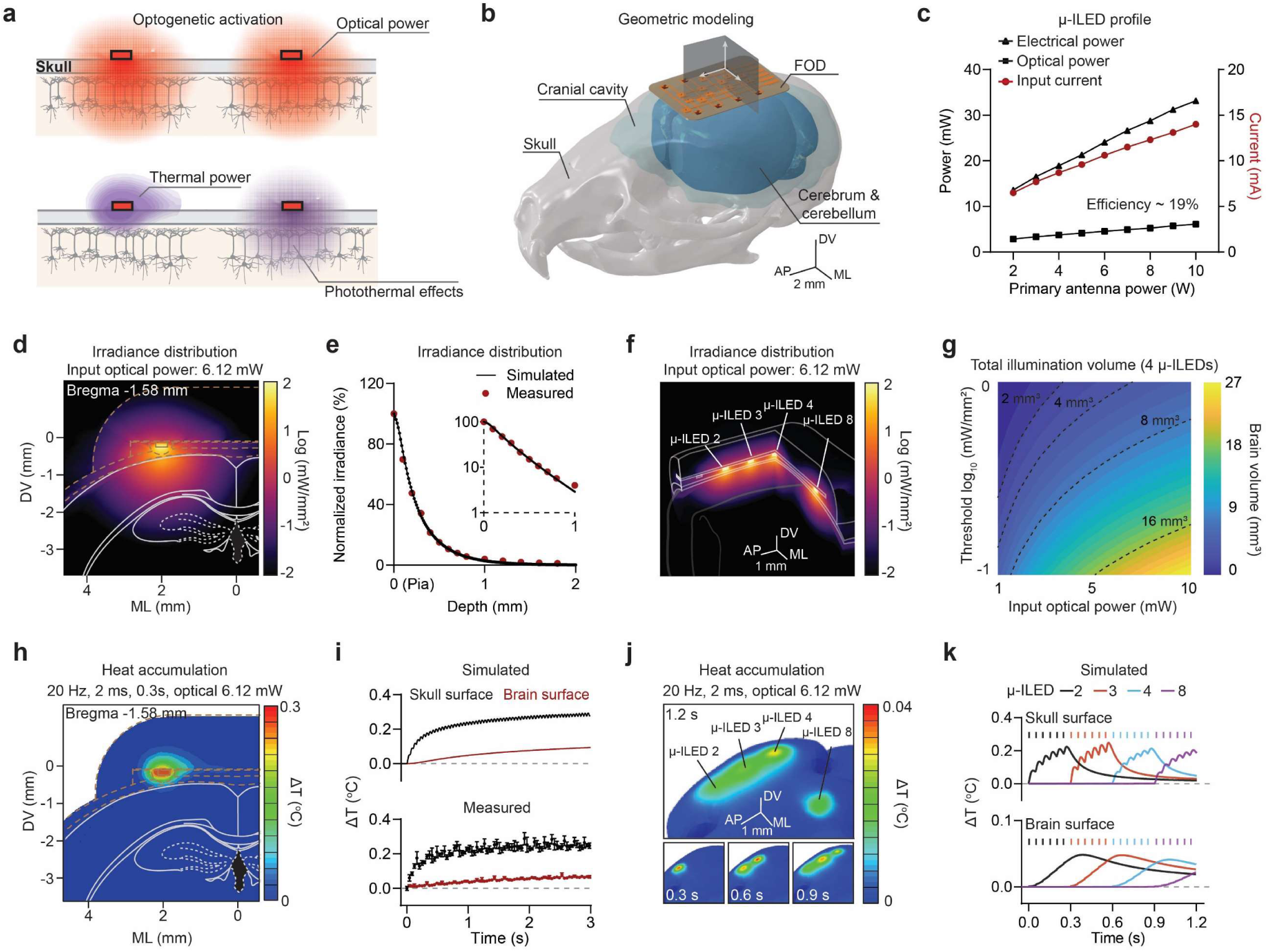
Spatiotemporal characteristics of light propagation and heat accumulation during device operation. (a) Power conversion pathways in the FOD, including the desired optical power for optogenetic activation, the non-optical power dissipating non-radiatively as heat, and the photothermal effects induced by tissue absorption of photons. Cartoon illustration, not reflecting actual scaling factors. (b) Geometric model of the main components surrounding the FOD. Accessory components, including dental cement, device encapsulations, and cyanoacrylate adhesive, are considered in the model but not displayed. Shading areas define the planes of interest in the simulations. (c) Electrical power, optical power, and input current for a single red μ-ILED (628 nm) as a function of primary power applied to the transmission antenna. (d) Coronal section of 3D Monte Carlo simulation of light propagation from a single μ-ILED relative to the brain regions defined by reference brain atlas at Bregma -1.58 mm. Input optical power of μ-ILED: 6.12 mW. (e) Summary data showing normalized simulated and measured optical intensity at different depths in the brain tissue. Absolute light intensity at 628 nm at different depths was normalized to the maximal intensity at pia (Z = 0). (f) Monte Carlo simulation of light propagation of 4-independent μ-ILEDs in three-dimensional space. Grey lines depict adjacent structures including dental cement, FOD, encapsulation, cyanoacrylate adhesive, skull, and brain surfaces. (g) Contour map plot showing the total illumination volume from 4 μ-ILED co-activation for different input irradiance and intensity thresholds for opsin variants. (h) Spatial distribution of heat accumulation during a single μ-ILED operation at 20 Hz (2 ms pulse width) for 0.3 s in the coronal section. (i) Simulated and measured temperature rise as a function of operation time at 20 Hz (2 ms pulse width) on the skull and brain surface. (j) Spatial distribution of heat generation on the brain surface with sequential 4 μ-ILED operation at 20 Hz (2 ms pulse width, 0.3 s each). (k) Simulated temperature rise as a function of operation time at 20 Hz (2 ms pulse width) on the skull and brain surface for 4 individual μ-ILEDs.

The effects of light and heat propagation can be captured by three-dimensional (3D) numerical simulations that use a comprehensive anatomy-guided model that includes the skull, cerebrospinal fluid, brain tissue, device materials, and surgical implantation materials (e.g. cyanoacrylate adhesive and dental cement) (**Fig. 2b**). A combination of constitutive properties of the materials and performance characteristics of the µ-ILEDs serve as inputs to this model (**Fig. 2c, Extended Data Fig. 5a-e, and Supplementary Tables 1-2**).

A coronal section of Monte Carlo simulation of the light penetration profile defines the irradiance distribution relative to the common coordinate framework for the mouse brain (**Fig. 2d**). Light intensity measurements at varying distances to a skull-mounted µ-ILED confirm the simulated attenuation profiles (**Fig. 2e and Extended Data Fig. 5f-g**). The results determine the depth and volume for optogenetic activation of ChrimsonR (628 nm; irradiance threshold of 1 mW/mm^2^) resulting from 6.12 mW optical power provided by a single µ-ILED^26^ (**Extended Data Fig. 5h**). Related simulations can define 3D light propagation profiles for the simultaneous operation of various combinations of multiple μ-ILEDs. As an example, **Fig. 2f-g and Extended Data Fig. 5i** define co-activation volumes and overlapping fractions from 4 µ-ILEDs (3 in the same column and 2 in the same row), for different optical power inputs and irradiance thresholds (0.1 to 1 mW/mm^2^). These approaches provide the basis for calibrating use of the system with different opsin variants.

Finite element analysis (FEA) techniques reveal transient heat generation and transport dynamics that provide guidelines for selecting operating parameters (power and duty cycle of the m-ILEDs) to ensure that the maximum temperature rise in the brain remains within a physiologically acceptable range. For optogenetic activation of excitatory opsin variants, a 2 ms pulse width is sufficient to generate action potentials in stimulated neurons with a 20 Hz frequency, corresponding to a 4% duty cycle. A 3D FEA thermal simulation based on these illumination parameters defines temperature distributions throughout surrounding structures for the operation of a single μ-ILED for 0.3 s (20 Hz, 2 ms, six pulses, parameter used in behavior experiments below). The results indicate a maximal temperature increase on the skull and brain surface at 0.24 and 0.04 °C, respectively (**Fig. 2h**). Temperature monitoring using customized thin-film thermistors confirms the simulated temperature increases (**Fig. 2i and Extended Data Fig. 6a- c**). The dental cement that surrounds the interface promotes the dissipation of heat due to its higher thermal conductivity (∼0.7 mW/mm/K, **Supplementary Table 2**) compared to that of the adjacent tissues (<0.6 mW/mm/K, **Supplementary Table 2**). The results also capture the time dynamics of the temperature distributions (**Fig. 2i**). An example in 3D space involves activation of multiple μ-ILEDs in sequence (4 μ-ILEDs, 0.3 s each for a total of 1.2 s, parameter used in behavior experiments below) to illustrate the spatiotemporal profile of heat generation across the regions of interest (**Fig. 2j**). At low input optical power (6.12 mW), thermal interference across multiple μ-ILEDs is minimal (**Fig. 2k, Supplementary Video 3**). **Extended Data Fig. 6d-g** shows the simulated and measured temperature rise on the skull and brain surface for a single μ-ILED operating with different duty cycles, frequency, and input optical power. Simple modifications to the device design, such as the addition of a metal coating on the backside of the FOD, can enhance thermal dissipation (**Extended Data Fig. 6h-i**), to further reduce the thermal load on the brain, as might be necessary for experimental protocols different from those reported here. Further details on numerical characterization and simulation of optical and thermal effects are included in the Materials and Methods section. The optical and thermal properties of materials are included in **Supplementary Tables 1-2**.

## Sequential cortical activation drives operant learning based on cue discrimination

To address the feasibility of generating artificial perception through digitized optogenetic inputs, we employed an operant conditioning behavioral paradigm, where animals learn to identify artificial cues and make associated actions to receive rewards^12,27^. Exploiting macroscale cortical gradients and their functional differentiation through spatial organization, we systematically manipulated neuronal activity in mouse cortical regions and quantitively measured perceptual responses in decision-making tasks.

We virally transduced AAV.hSyn.ChrimsonR.tdT into a total of eight cortical regions, spanning four distinct areas in both hemispheres, including the motor cortex (AP: +1.5 mm, ML: ±2.0 mm), somatosensory cortex spanning limb regions (AP: 0 mm, ML: ±2.0 mm), somatosensory cortex covering trunk representation (AP: -1.5 mm, ML: ±2.0 mm), and visual cortex (AP: -3.0 mm, ML: ±2.0 mm) (**Extended Data Fig. 7a-d**). The efficacy of transcranial optogenetic activation was visually confirmed by tonic motor responses elicited by prolonged high-frequency (20 Hz) light stimulation at the motor cortex, as described in our previous reports^21,28^.

In the operant conditioning paradigm, mice initiated trials by poking the center port. Subsequently, sequential cortical optogenetic inputs were delivered to four designated locations among the eight, each stimulated for 0.3 s at 20 Hz (6 pulses with 2 ms pulse width). Following stimulation termination, mice received a “go” cue indicating they could choose the left or the right-side port. If mice selected the port associated with the stimulation pattern, they received a sucrose reward droplet, while wrong choices led to an air puff punishment (**Fig. 3a**).

**Fig. 3.**
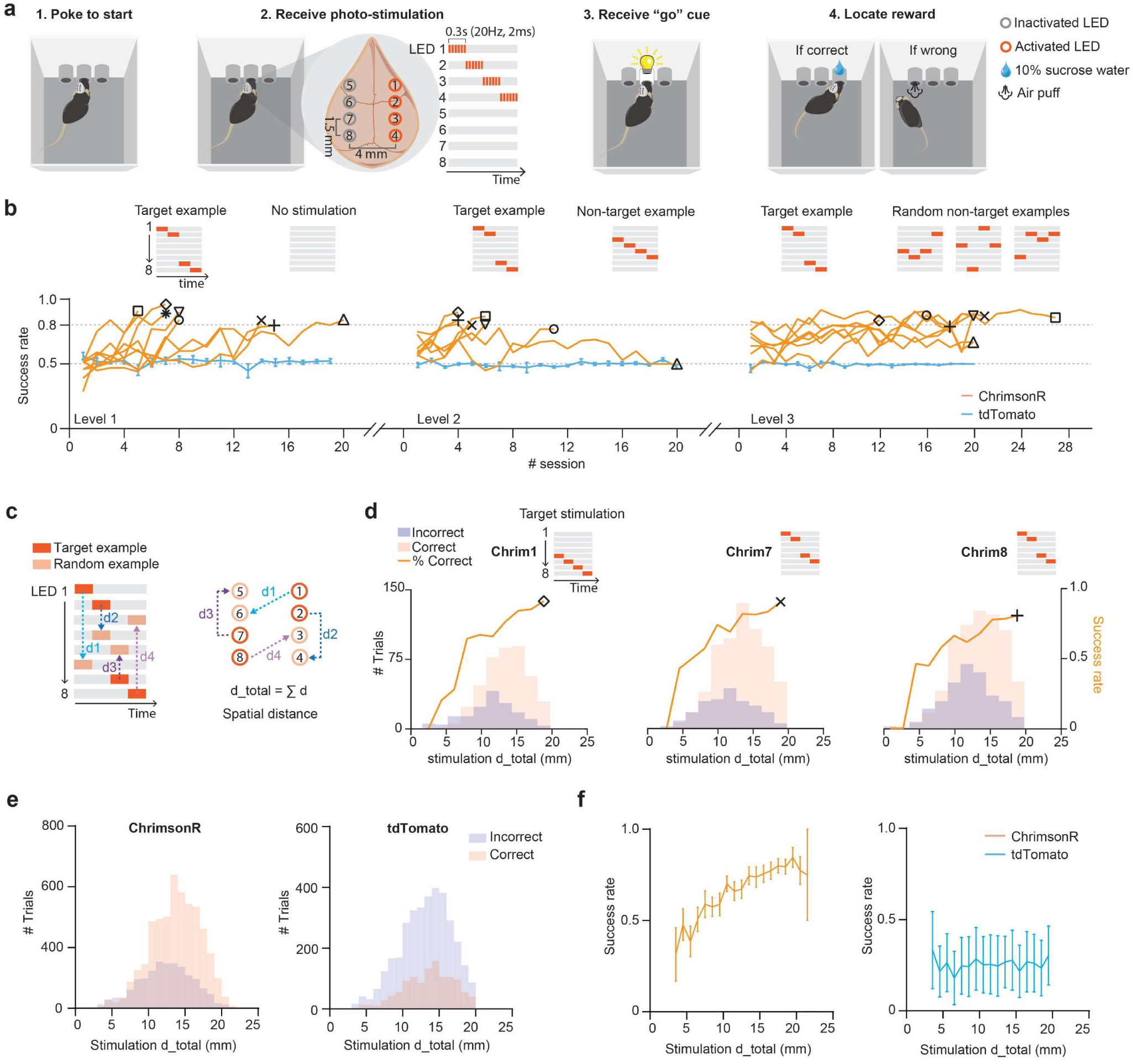
Sequential cortical activation drives operant learning based on cue discrimination. (a) Schematic illustration of the operant learning task with artificial optogenetic cues. Mice make decisions based on the discrimination of cortical activation sequences, with successful trials rewarded by sucrose. (b) Summary data showing the performance trajectories of cue discrimination for animals expressing ChrimsonR or tdTomato based on the success rate of obtaining sucrose rewards. Level 1, distinction of target sequence versus no stimulation; Level 2, target sequence versus a certain non-target sequence; Level 3, target sequence versus a pool of randomized sequences. Chance = 0.5. n = 7 - 8 animals for ChrimsonR and 5 animals for tdTomato. Each session includes up to 100 trials. Shapes indicate individual animals. (c) Definition of spatial distance between the target sequence and individual randomized non-target sequence in Level 3. (d) Line plot showing success rate as a function of spatial distance for 3 individual animals in the Level 3 task. The number of trials with correct or incorrect choices in each bin of spatial distance was plotted in the histogram. (e) Summary data showing the number of trials with correct or incorrect choices in each bin of spatial distance from all animals expressing ChrimsonR or tdTomato. n = 7 animals for ChrimsonR and 5 for tdTomato. (f) Summary data showing success rate as a function of spatial distance for all animals expressing ChrimsonR or tdTomato. n = 7 animals for ChrimsonR and 5 for tdTomato.

We trained mice to discriminate a designated target sequence in three tasks of progressive difficulty. The simplest task required discrimination against null-stimulation, where none of the LEDs lit up. After achieving a high success rate (80%), mice had to discriminate the target sequence against a specific non-target sequence. After the pretraining at level 1 and 2, mice received a pool of randomly generated non-target sequences in the trials, requiring discrimination of the target sequence from this pool (**Fig. 3b**). ChrimsonR-expressing mice learned to reach higher success rates of getting rewards in the three levels of tasks within 10 days (median number of sessions: level 1, 8; level 2, 5; level 3, 14.) (**Extended Data Fig. 7e-f**), while the performance of tdT-expressing control mice maintained at chance level (0.5) after 20 sessions in each task level (**Fig. 3b**). ChrimsonR-expressing mice exhibited a performance increase beyond chance level, demonstrating robust perception from digitized optogenetic inputs as a driver of decision-making processes for reward localization.

Because of non-uniform connectivity across the broader cortical space^29–31^, it is unknown whether a larger physical distance between stimulation locations correlates strongly with stimulus discriminability. We therefore utilized the randomized non-target sequences from the level 3 task to construct a spatial distance framework correlating sequence discrimination performance with spatial distance on the cortical map (**Fig. 3c**). Distinct distributions of success and failure trials based on the spatial distance of the randomized sequence were observed in the data collected from individual animals. The success rate increased as the non-target sequence spatial distance from the designated target sequence increased (**Fig. 3d**). The distribution of trial outcomes and the trends in success rate persisted at the group level. In contrast, trial outcomes from tdT-expressing control animals were not associated with the spatial distance of sequences (**Fig. 3e and 3f**). These findings suggest the spatial distance among sequential cortical activations may represent a ’cognitive distance’ for discrimination. Control analyses on reaction time distribution and individual animal behavioral analysis are detailed in **Extended Data Fig. 8a-b, and 9.**

## Artificial cortical inputs follow the primacy principle of perception

To investigate whether perception rules extend to artificially generated cortical activation patterns, we treated the digits of the sequence as independent variables. Randomly generated non-target patterns were categorized based on stimulated cortical locations for each digit. For example, non-target stimulation trials initiated with motor cortex stimulation were grouped, and the total success rate for these trials was calculated (**Fig. 4a**). Hypotheses, rooted in two principles of perceptual psychology, were applied to this framework, considering the contribution of each digit to perception discrimination based on their temporal sequence. The primacy hypothesis^32^ posits that the earlier stimulated cortical region should contribute more to perceptual discrimination, compared to later stimulated regions **(Fig. 4a**). In contrast, the recency hypothesis^33^ suggests that the later incoming information is weighted more heavily in decision-making processes. Therefore, the primacy hypothesis predicts that performance will be worse when there is overlap between target and non-target stimuli early in the sequence, while the recency hypothesis suggests that performance will be worse when there is overlap late in the sequence.

**Fig. 4.**
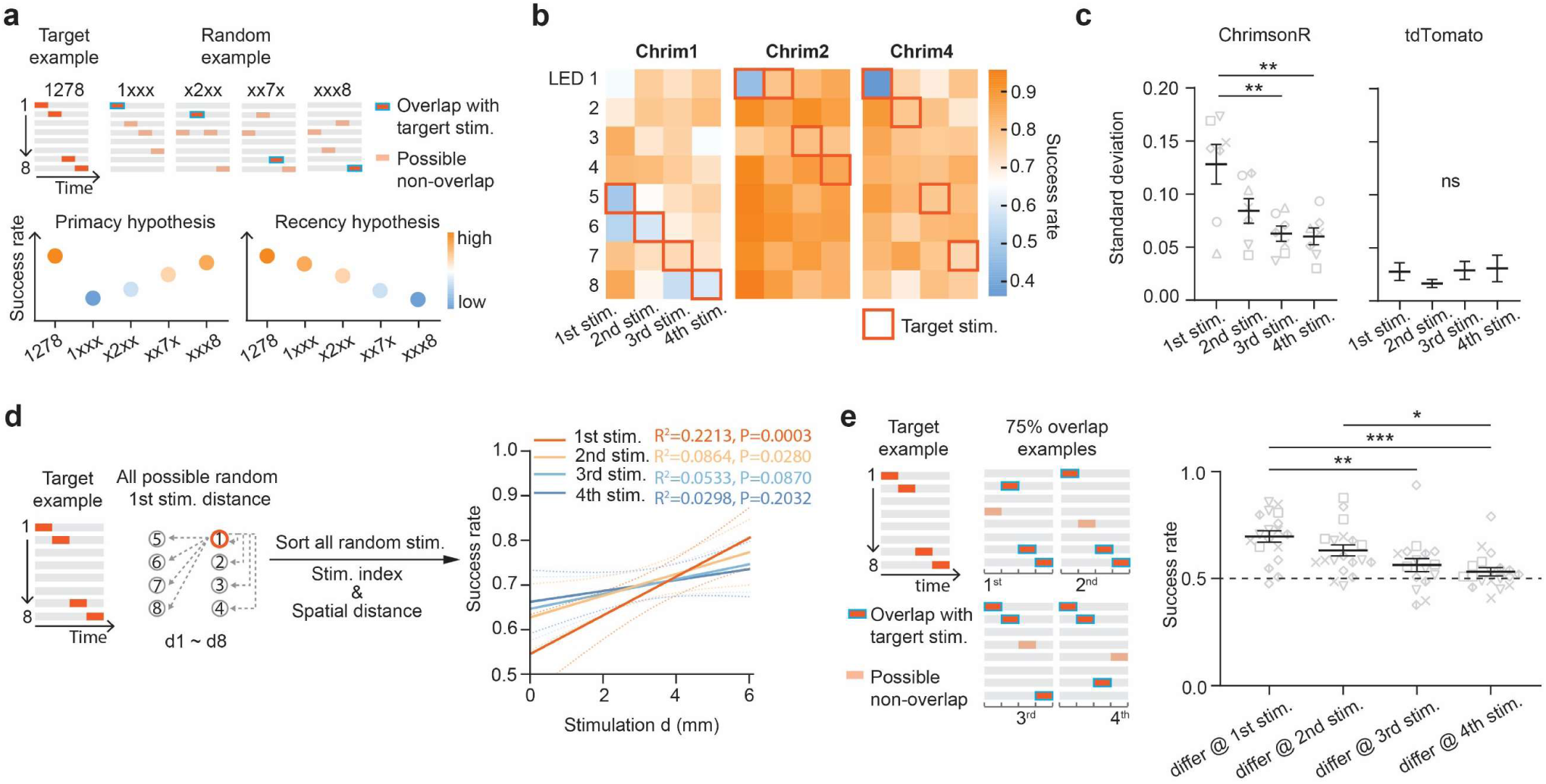
Artificial percepts follow the primacy principle of perception. (a) Hypothesized outcomes in the discrimination of sequential cortical activation based on the primacy and recency principles of perception. (b) Heatmap showing success rate for randomized non-target sequences grouped by specific stimulation locations at the first to the fourth stimulation digit in three ChrimsonR expressing animals. Red squares indicate the target stimulation sequence for each animal. (c) Summary data showing the standard deviation of success rates from the eight possible stimulation locations in the individual stimulation digits for ChrimsonR expressing animals and tdTomato controls. One-way ANOVA, Sidak’s multiple comparisons, ChrimsonR, p = 0.0020, F (3, 24) = 6.657, 1st stim vs 3rd stim: p = 0.0046, 1st stim vs 4th stim: p = 0.0032; tdTomato, p = 0.6745, F (3, 16) = 0.5200; n = 7 animals for ChrimsonR and 5 for tdTomato. (d) Left, schematic illustration of non-target stimulation sorting based on the spatial distance from the target stimulation at each stimulation digit. Right, Pearson’s correlation analysis of spatial distance and success rate based on stimulation digit. (e) Experimental validation of the primacy principle of perception. Left, schematic illustration of experimental design. A group of probing sequences with 75% similarity (3 stimulation digits) to the target sequences were used as non-target stimulations. Varying stimulation locations were selected at each non-overlapping digit for each animal. Right, summary data showing success rate in the 75% similarity probing tasks for each non-overlapping digit. Each data point represents a single 100-trial session, and each shape represents the animal used in Figures 3-4. One-way ANOVA, Sidak’s multiple comparisons, p < 0.0001, F (3, 69) = 8.327, differ @ 1st stim vs differ @ 3rd stim, p = 0.0030, differ @ 1st stim vs differ @ 4th stim, p = 0.0001, differ @ 2nd stim vs differ @ 4th stim, p = 0.0308; n = 5 animals.

With specific parameters tuned in this operant conditioning paradigm, our results demonstrated that overlapping stimulated cortical regions between target and non-target sequences at the first digit resulted in the lowest success rate, supporting the primacy hypothesis (**Fig. 4b, Extended Data Fig. 10a**). Furthermore, switching the stimulation region at the first digit resulted in higher variation in the performance of cue discrimination (**Fig. 4c**). As performance varies the most with the identity of the first digit of the non-target stimulus, it contributes maximally to perceptual discrimination. Additionally, spatial distance effects on perceptual discrimination were differentially modulated by the temporal sequence. Adding spatial distance to the framework revealed that the spatial distance between stimulated regions at the first digit correlates most strongly with behavioral outcomes (**Fig. 4d, Extended Data Fig. 10b-c**).

To further explore our observations consistent with the primacy principle, we designed a similarity probing task. After the three levels of learning, mice were presented with a group of probing non-target sequences that were similar to the target sequences. In this task, mice had to distinguish between the target sequences and a non-target sequence with 50% to 75% similarity (2-3 overlapping digits). Mice performed significantly better in distinguishing 50% similarity sequences compared to 75% (**Extended Data Fig. 10d-e**). In the 75% similarity probing task, we further tested the primacy hypothesis by switching the non-overlapping stimulation across the four digits (**Fig. 4e**). Mice distinguished the 75% similar non- targeting sequence around chance level when the overlapping stimulation involved the first 3 digits. They performed significantly better in the task where the overlapping stimulation occurred at the last 3 digits. These results demonstrate that the artificially generated cortical inputs can be tuned to follow some of the principles (e.g., similarity and primacy) established in the literature on perception^34^.

## Discussion

In this study, we developed a fully implantable optogenetic neural interface, and its supporting ecosystem of technologies, for wireless transmission of digitized information to a large-scale network of neurons. The reported device utilizes an array of individually addressable optical stimulators to transcranially deliver regional programmable spatiotemporal patterns of illumination to targeted cortical areas. Real- time programmability provides multiple degrees of freedom, enabling the delivery of dynamic perceptual feedback signals directly to the cortex through patterns of multi-channel optogenetic stimulation. The mechanically compliant fully subdermal-implantable form factor allows application in freely moving animals without external physical constraints, facilitating the study of artificial perception delivery within behavioral contexts. Furthermore, comprehensive numerical simulations on through-the-skull light penetration and thermal dissipation profiles serve as detailed guidance for future technology designs for relevant neuroscience applications. We expect that this powerful and scalable platform is generalizable to accommodate various animal models.

Utilizing this technology, we constructed a framework to investigate features of artificially synthesized cortical activation sequences in the perceptual domain. We measured discrimination-based perceptual responses to systematic input manipulations—a fundamental approach in psychology applied to various classes of external stimuli to reveal the principles underlying perception^35^. Our continued advancements in wireless optical techniques allow causal manipulation of the spatiotemporal neural representation at large-scale cortical levels. We observed that perceptual responses correspond to the similarity in spatial features, which are further modulated by the temporal order of cortical activation.

Our primary results demonstrated a serial-position effect modulating the similarity among artificially synthesized perceptions. Notably, the adaptability of the platform, coupled with open-source tools in the neuroscience community, holds the promise of addressing different fundamental rules of perception that may be applied in brain-machine interface development, further expanding the dimensionality of the artificial neural syntax that can be perceived by the brain.

Future development of this device platform involves enhancing the resolution of addressed neuron populations by increasing the number and reducing the size of optogenetic stimulators. One promising route is to introduce digital control methods to address multiple optoelectronic components with reduced wiring^36,37^. Additionally, transferring the optogenetic stimulator array into a compliant substrate will further enable the applications in peripheral nervous systems and other organs^38^, promoting innovative advancements in neural interface technology.

## Methods

### Device fabrication and assembly

A third-party vendor (PCBWay) produced the flexible printed circuit boards (fPCB) via laser ablation for the electronic module, the µ-ILED panel, and the serpentine interconnect based on designs created using commercial CAD software (AutoCAD, Autodesk). The fPCB substrate for the electronic module and µ- ILED panel used a three-layer copper (18 µm) - polyimide (25 µm) - copper (18 µm) fPCB design. Alternatively, polyester (PET, 64 µm) may be used to replace polyimide to enhance the optical transparency of the devices. The fPCB substrate for the serpentine interconnect used a five-layer copper (12 µm) - polyimide (12.5 µm) - copper (12 µm) - polyimide (12.5 µm) - copper (12 µm) fPCB design. The total thickness was ∼100/200 µm for polyimide/PET-based circuit boards with solder mask layers, and ∼60 µm for serpentine interconnects. Hot air soldering using a heat gun operated at 220-270 °C and low-temperature solder paste (Indium Corporation) electrically bonded the commercial electronic and optical components on the fPCB to establish the function of the electronic module and the µ-ILED panel. The same soldering strategy further electrically connected the electronic module and the µ-ILED panel with the serpentine trace to form the complete device. The chemical vapor deposition (CVD) process then formed a uniform parylene encapsulation coating (14 μm) around the device. Additional dip-coated epoxy (Norland 63) cured by ultraviolet (UV) light exposure mechanically secured and encapsulated the solder joints. Finally, mold-cast silicone coatings (Ecoflex 00-30, 400 μm) symmetrically covered the top and bottom surfaces of the serpentine trace and electronic module.

### Smart NFC wireless electronic device

A 10-turn planar coil surrounding the perimeter of the device enabled magnetic inductive coupling to an external antenna for near-field communications (NFC) and wireless power transfer. The harvested power was buffered on a five-22 µF ceramic capacitor bank that allowed storage of ∼1.3 mJ of charge when completely charged (5 V). The harvested power was regulated with a 2.8 V low drop-out regulator to provide constant voltage to the electronics. The NFC memory (M24LR04E-R, STMicroelectronics) provided standardized (ISO 15693) NFC communication with an external RF NFC driver (PDC box, Neurolux, Inc.) as the basis for bidirectional communication. A low-power 8-bit microcontroller (Attiny84, Microchip Technology, Inc.) equipped with dedicated firmware received remote commands through the NFC memory and implemented spatiotemporal activation of the eight µ-ILEDs (TCE12-628, Three-five materials).

### Hybrid analog/digital matrix multiplexing

The independent control of the eight µ-ILEDs is based on matrix multiplexing where two variable voltage lines connect the rows at the anodes of the µ-ILEDs and four digital lines connect to the columns at the cathodes (**Extended Data Fig. 4**). The use of two independently addressable digital-to-analog converters (DACs) connected with the microcontroller via I2C provides user-defined voltage levels. The use of the other four digital control lines allows for forward biasing only when writing logic 0 but not otherwise. This combination of hybrid analog/digital multiplexing serves as the basis for independent µ-ILED activation at programmable voltages.

This simple matrix intensity modulation approach requires only 2 DAC chips (2 mm × 2 mm each, MCP4706, Microchip Technology Inc.). In contrast, other intensity control implementations increase the component count and complexity of the electronics, posing an integration challenge in the limited space available in the implantable device. These alternative approaches include the Howland voltage-controlled current source (one operational amplifier chip and four passive resistive elements) and a voltage-controlled transconductance amplifier current source (one operational amplifier chip, one transistor, one resistor), where both cases also require the use of a voltage source such as the DACs.

### Implementation of wirelessly programmable, intensity-controlled spatiotemporal illumination

The wireless device generates programable spatiotemporal patterns of activation using the following approach. A read/write RF-accessible non-volatile memory (M24LR04E-R, STMicroelectronics) receives commands via NFC. This NFC memory, connected with the microcontroller (Attiny84, Microchip Technology, Inc.) via I^2^C, produces an interruption to the protocol upon command arrival. The microcontroller, prompted by this interruption, reads a memory block consisting of four bytes that contain the sequence/activation order for the four µ-ILEDs. Each byte includes the information for the µ-ILED index to activate (4 least significant bits) and up to sixteen intensity levels (4 most significant bits). The channel selection and intensity are then sent to the analog/digital hybrid matrix multiplexer to forward bias to the selected µ-ILED at one of the sixteen preprogrammed voltage levels. This block of memory is written via NFC on demand as required by the progression of the behavioral protocol. It is worth mentioning that the sixteen preprogrammed voltages can be updated via NFC before the start of the experiment. Operational parameters such as frequency and pulse widths per µ-ILED and delay between µ-ILEDs in the sequence are also user-defined via NFC. These values, once written, will remain in memory until further update, if necessary. This way, universal firmware can be used without the need for firmware customization per application.

### Mechanical modeling and characterization

The commercial software Abaqus (2020, Dassault Systemes Simulia Corp.) was used to assess the mechanical characteristics of the device. The circuit board, the FOD, and the serpentine interconnect were modeled by composite shell elements (S4 elements), consisting of films of copper, polyimide and parylene. The Young’s modulus and Poisson’s ratio of the film materials are: ECu = 130 GPa, νCu = 0.34, EPI = 2.5 GPa, νPI = 0.34, EParylene = 2.1 GPa, νParylene = 0.34 ^19^. All components were embedded in silicone (Ecoflex 00-30) encapsulation. The encapsulation was modeled using tetrahedron elements (C3D10H). Mooney-Rivlin hyperelastic constitutive model was employed to characterize the encapsulation material, with CEcoflex10 = 8 kPa, CEcoflex01 = 2 kPa, DEcoflex = 0.002 kPa^-1^ (corresponding to initial Young’s modulus EEcoflex = 60 kPa and Poisson’s ratio νEcoflex = 0.49) ^39^. For stretching, twisting, and bending simulation, the circuit board was fixed and the displacements and rotations were applied to the panel. For the contact simulation of the panel to the skull, a uniform pressure was applied to the panel until all µ-ILEDs contacted the skull. The element number in the model was ∼6 × 10^5^, and the minimal element size was about 1/4 of the width of the narrowest copper trace (12 µm). Geometric nonlinearities were considered in the simulation. The finite element mesh convergence of the simulation was validated for all cases.

### Electromagnetic modeling and power characterization

To confirm the magnetic field within the cages was substantial and uniform for the behavior experiments, the software COMSOL (COMSOL 6.0) was used for the characterization of the double-loop antenna configuration. An adaptive mesh composed of tetrahedron elements and a spherical surface with a radius of 1,000 mm were employed as the radiation boundary to ensure computational precision. For the numerical modeling, the relative dielectric constant of air was set to 1, and the conductivity of copper was set to 5.8 × 10^7^ S m^−1^. To evaluate the normalized magnetic induction intensity on different planes of the cage, the current flowing through the copper wires was set to 1 A.

### Optical characterization

The optoelectronic characterization of µ-ILED began by obtaining the corresponding I-V characteristics using a semiconductor device analyzer (Keysight S1500A) interfaced with a probe station (Signatone 1160). Then, measurements of µ-ILED current during wireless operation, combined with obtained I-V characteristics, yielded the electrical power of µ-ILED with respect to the power supplied to the transmission antenna. An integrating sphere photodiode power meter (S140C, Thorlabs Inc. Newton, NJ) in combination with a calibrated power meter (PM100D, Thorlabs Inc.) served to characterize the optical power generated by the µ-ILED. A current source (Keithley 6221, Tektronix Inc., Beaverton, OR) powered the µ-ILED after insertion into the integrating sphere, with applied current from 1 mA to 20 mA with a 1 mA interval. This result, combined with µ-ILED I-V characteristics and current during operation, yielded the µ-ILED efficiency and output intensity with respect to the transmission antenna power.

For measurements involving light attenuation in brain tissue, a patch cable (600 µm core, NA 0.37, Doric, Québec, CA) attached to an optical fiber (600 μm core, NA 0.37, 6-mm long) relayed the signal to the integrating sphere for recording the intensity. The isolated mouse skull and brain tissues, affixed to the µ-ILED array, were positioned on a metal block on a stereotaxis that also secured the optical fiber. With 0.1 mm precision, the optical fiber advanced through the skull base and brain, and light intensity readings were recorded at each depth until the fiber tip reached the pia. The absolute intensity readout was normalized to the maximum intensity at the pia level, providing the light attenuation ratio in the brain tissue at each depth. Subsequent measurements of post-skull optical penetration efficiency followed the same procedures, incorporating an additional piece of mouse skull affixed to the µ-ILED exposure surface when utilizing the integrating sphere for measurements of optical power for the µ-ILED with varying inputs.

### Thermistor fabrication and thermal characterization

To measure the temperature load in the surrounding structures of the optical-neural interface, a thin-film thermistor was fabricated using standard photolithography. Fabrication of the thin-film thermistor began by spin-coating polymethyl methacrylate (PMMA; 500 nm) and polyimide (PI; 5 μm) on a silicon wafer substrate sequentially. Electron beam physical vapor deposition (EBPVD) then formed a thin conductive film of 20 nm Ti and 100 nm Au on the PI layer. Another spin-coating process formed a thin layer of photoresist (Microposit S1813, ∼1.5 μm) on the metal layers. A maskless aligner (Heidelberg MLA 150) patterned the photoresist layer into the desired geometry that contains sensors and connection traces. Sequential wet etching using Au etchant (Transene, Type TFA) and buffered oxide etch (BOE 6:1; Avantor) transferred the photoresist pattern to the underneath metal layers. After removing the remaining photoresist with acetone, a second layer of PI (5 μm) and a thin layer of Cu (60 nm) were deposited sequentially using spin-coating and EBPVD, respectively. A similar lithography and wet etching process (using Cu etchant) formed an etching mask on the Cu layer that encapsulated the patterns on the Ti/Au layers. A dry etching process using reactive ion etching (March RIE) transferred the Cu layer pattern to the underneath PI. Finally, the Cu mask was removed by Cu etchant.

Two clamps firmly held another same-size silicon wafer against the device silicon wafer with a tissue sandwiched in between. The wafers were then placed in an acetone bath in a glass beaker. A hotplate operated at 80 °C warmed the bath to accelerate the dissolution of PMMA. After 2 hrs, a water-soluble tape transferred the device from the silicon wafer after careful removal of the top wafer and tissue. The thermistor was wired to a digital multimeter (USB-4065, National Instruments, Austin, TX) after the removal of the backside water-soluble tape with deionized water. The temperature responses of the sensors underwent calibration using a data logger thermometer (HH374, OMEGA, Norwalk, CT). The thin-film thermistor was then attached to the surface of the μ-ILED, skull, and brain with epoxy glue to acquire temperature changes during µ-ILED operation.

### Geometric modeling of cranial structures

The geometric models of the cranial structures were generated from an open-source database of rodent anatomical structures, corrected for C57/BL6 mice based on species-specific dimensional profiles. Original files (.stl) were converted to .stp and modeled using Abaqus. The FOD was reconstructed from the CAD file, mounting on the surface of modeled cranial structures. Additional materials, including cyanoacrylate adhesives, Norland 63, parylene-C, µ-ILEDs, fPCB, and dental cement, are modeled in the full model. The geometric model was further converted into voxels and finite elements for optical and thermal simulation.

### Optical simulations

The Monte Carlo method was used to simulate the optical profile for transcranial optogenetic stimulation^40^. Each volume in the numerical simulations consisted of (400)^3^ voxels, with dimensions of (12.5 µm)^3^, filling a total volume of (5 mm)^3^. The red (628 nm) illumination source corresponding to the dimensions of the µ-ILED was set to 0.3 × 0.3 × 0.09 mm^3^, featuring a Lambertian emission angle distribution:

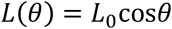

*L*(*θ*) is the optical intensity. *θ* is the angle between the irradiation direction and the normal of the µ-ILED lower surface, which is 0.8 mm above the center of the numerical simulation volume. *L*0 is the optical intensity for *θ* = 0. The simulation launched 10^9^ photons. The materials’ optical properties, including absorption coefficients (*μ*a), reduced scattering coefficients (*μ’*s), and refractive index (*n*) are summarized in **Supplementary Table 1** for 628 nm. After performing the simulation for each volume of µ-ILED, postprocessing of the optical fluence rate yielded illumination profiles in three-dimensional space. The three-dimensional reconstruction of the transcranial illumination profiles resulting from 6.12 mW input optical power was generated using Paraview 5.7.0 (Kitware, Clifton Park, NY). The volume of µ-ILEDs co-illumination and crosstalk was then calculated based on input optical power and irradiance threshold for opsin excitation.

### Thermal finite element analysis model

The transient-state thermal propagation considering the thermal load of the µ-ILEDs and the body heat flux caused by the optical power is defined by the governing equation ^19^:

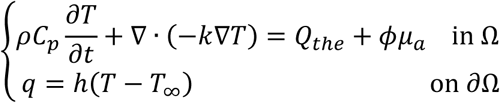

Here, *T* is the temperature, *ρ* is the mass density, *Cp* is the heat capacity, and *k* is the thermal conductivity. *Qthe* is the heat source generated by the thermal load of the µ-ILEDs. *ϕ* is the optical fluence rate calculated in the optical simulation. Ω is the model region. The convective heat transfer boundary conditions are assigned on the boundary of this model region (∂Ω). *q* is the surface heat flux. *T*∞ is the environmental temperature, and *h* is the convective heat transfer coefficient. Here, *h* = 0.025 mW/mm^2^/K for the dental cement boundary. *h* = 0.005 mW/mm^2^/K for the skull boundary. Abaqus was utilized to compute the governing equation using the finite element method. Linear tetrahedral element was used, and the minimum edge length was 60 μm. The thermal conductivity (*k*), heat capacity (*Cp*), and mass density (*ρ*) of the materials are summarized in **Supplementary Table 2**. Additional measurements using a previously reported thermal actuator and multisensory (TAS) module ^41^ confirmed the thermal conductivity of essential biological and device materials.

### Scanning electron microscopy (SEM)

An FEI Quanta 650 environmental scanning electron microscope examined the surface morphology of the µ-ILED array, and serpentine interconnects in high-vacuum (HV) secondary electron (SE) mode. An electron beam of 10 kV and a spot size of 3.0 were used. The surfaces of the samples were directly exposed to the electron beam during characterization.

### Animals

Animals were handled according to protocols approved by the Northwestern University Animal Care and Use Committee. C57BL/6 mice were obtained from Charles River (Wilmington, MA) and bred in-house. Adult male and female mice, aged between postnatal days 80 to 120 and weighing 20 to 30 g at the onset of experiments, were used in the study. Prior to procedures, all mice were group housed, kept at approximately 25°C, and maintained on a standard diet under a 12-hour light and 12-hour dark cycle (lights on at 6:00 or 7:00). A plastic igloo shelter and nesting materials were used as environmental enrichment. Littermates were randomly assigned to conditions. Mice were not involved in any previous procedures and only healthy and immunocompetent mice were used.

### Stereotactic injections and device implantation

Mice were anesthetized with isoflurane (3% for induction, 1.5-2% for maintenance) and fixed on a small animal stereotaxic frame (David Kopf Instruments, Tujunga, CA) for intracranial injections. Analgesia included bupivacaine, meloxicam, and buprenorphine-ER. AAV1.Syn-ChrimsonR-tdT (1 × 10^13^ GC/mL, Addgene viral prep #59171-AAV1, a gift from Dr. Edward Boyden), or AAV8.FLEX.tdT (1 × 10^13^ GC/mL, Addgene viral prep #28306-AAV8, a gift from Dr. Edward Boyden) mixed with AAV1.hSyn.Cre.WPRE.hGH (1 × 10^12^ GC/mL, UPenn viral core, a gift from Dr. James M. Wilson, unpublished) was delivered through a pulled glass pipette (Tip diameter 7 - 10 µm) for a total volume of 200 nL at a rate of 100-150 nL/min using an UltraMicroPump (World Precision Instruments, Sarasota, FL). Injection coordinates for the motor cortex, primary somatosensory cortex (limbs), primary somatosensory cortex (trunk), and primary visual cortex are AP: +1.5, 0.0, -1.5, and -3.0 mm; ML: ± 2.0 mm; DV: -0.5 mm. Viral vectors were expressed for 2-4 weeks before device implantation.

Device implantation followed the procedure for back subdermal implantation reported previously^42^. Initially, an incision was made along the midline of the scalp to expose the skull. The FOD was positioned on the skull, aligning individual LEDs with the pre-made burred holes for viral injections. Subsequently, another 1 cm incision was made at the midline of the back, approximately at the T10-L1 level. The subcutaneous fascia between the scalp and back incisions was separated, and the wireless device was gently pulled from the scalp incision towards the lumbar spine. Serpentine interconnects linking the FOD and the electronic board were routed subcutaneously through the neck. Following this, the incisions were sutured closed, and the animals were closely monitored and allowed to recover for several hours before being returned to the colony for appropriate post-surgical observation.

### Tissue processing and histology

Mice were deeply anesthetized using isoflurane and subsequently underwent transcardial perfusion with 4% paraformaldehyde (PFA) dissolved in 0.1 M phosphate-buffered saline (PBS). Following perfusion, brains were post-fixed in 4% PFA at 4 °C and then rinsed in PBS before being sectioned at a thickness of 60 µm using a vibratome (Leica Biosystems). These sections were then mounted onto Superfrost Plus slides (Thermo Fisher Scientific), allowed to air dry, and coverslipped using a mixture of glycerol and TBS in a 9:1 ratio, with Hoechst 33342 at a concentration of 2.5 μg/mL (Thermo Fisher Scientific). Sections were scanned using an Olympus VS120 slide scanning microscope (Olympus Scientific Solutions Americas, Waltham, MA) to assess the areas of viral expression.

### Experimental setup for behavioral studies

A host computer running a customized MATLAB (MathWorks) script serves as the integrator of our wireless device with the open-source mouse operant chamber (Bpod 1030, Sanworks LLC, NY, developed by Josh Sanders). The system integration is as follows. The Bpod system comes with a dedicated library that interfaces with MATLAB, providing access to the sensors and actuators contained in the hardware of the operant chamber. For our wireless device, we developed a MATLAB library that communicates with the NFC RF module contained within the power distribution and control box (PDC Box, Neurolux Inc) using a serial interface (RS232-to-USB converter). Using this library and the NFC reader/writer we gained real-time control of the operation parameters of the NFC optogenetic device such as programmable μ-ILED stimulation patterns and intensity, frequency, pulse widths, and RF power of the PDC Box. The PDC Box drives a dual-loop antenna that wraps around the Bpod mouse operant chamber. Impedance and loading matching between the RF power module and the homemade RF antenna were controlled by an antenna tuner (Neurolux) to ensure efficient power transmission and signal communication. The customized MATLAB scripts and the multisystem integration of technologies allowed to control the state machine of the Bpod to implement real-time adjustment of stimulation parameters based on behavioral feedback.

### Water restriction

Mice started water restriction 5 days before training to motivate reward-seeking behaviors. The procedure began with recording initial body weight. Mice received 45% of the free-drinking water requirement (2.8 mL/30 g, daily) per Jackson Labs standards. The daily water supplement was calculated as 0.45 × 2.8 × (original body weight) / 30 g. Mice were weighed and monitored daily for health status. If body weight dropped below 85%, additional water was provided (0.85 × original weight - current body weight). Water was dispensed in a plastic Petri dish in each home cage. During behavioral experiments, the total daily water included both the reward from the Bpod operant chamber, and the extra amount given after sessions. Mice received their full calculated water requirement in their home cage, if they did not engage in any reward-seeking behaviors ^43^.

### Operant conditioning

The operant conditioning paradigm involved two phases: habituation-pretraining before device implantation and stimulation training after implantation. During pretraining, mice were acclimatized to the operant chamber and trained to follow the trial structure. Each trial started when the mouse poked the center port, and a 10% sucrose water reward was provided after a second poke at either the left or right port. A white light at the center port served as a “go cue”. As mice became more engaged in the task, the required holding time at the center port gradually increased to 1.0 s to initiate a trial. Once animals completed 100 trials within 30 min, they proceeded to the training phase following device implantation. In the stimulation-training sessions, poking the center port triggered a sequential red light stimulation pattern from the µ-ILED array, consisting of four stimulation locations (20 Hz, 2 ms pulse width, 0.3 s at each location), with no inter-stimulus interval. The reward port remained unresponsive to premature pokes until the stimulation sequence was complete. Following the stimulation sequence, a “go-cue” was given at the center port, and the animals were allowed to locate rewards. Initially, each mouse was assigned a “target stimulation” pattern that remained constant throughout training, always indicating a reward on the left side. Incorrect choices resulted in an air puff punishment (<10 psi, 0.2 s). Mice advanced to the next task level upon achieving an 80% success rate or completing 2,000 trials at the current level. No left/right port bias correction methods were employed. In all pretraining and training sessions, an external red-light source was used to mask non-specific optical inputs from the environment and the FOD.

### Microscale computed tomography (microCT) imaging

An animal implanted with the wireless optogenetic encoder was euthanized and imaged with a preclinical microCT imaging system (Mediso nanoScan scanner). Data acquisition parameters include medium magnification, 33-μm focal spot, 1 × 1 binning, and 720 projection views (300-ms exposure time) over a full circle. Reconstruction of the projection data utilized a voxel size of 68 μm and post-processed with Mediso Nucline (version 2.01).

### Statistical analyses

Required sample sizes were estimated based on previous publications and experience ^44–46^. At least two internal replications are present in the study. The number of biological replicates, trials, and animals is reported in Figure legends. Animals were randomly assigned to treatment groups; no data were excluded for analyses. Group statistical analyses were done using GraphPad Prism software (GraphPad, LaJolla, CA). All data are expressed as mean ± SEM, box-whisker plot (minimum, first quartile, median, third quartile, and maximum), or individual plots. Statistical significance was determined by two-tailed Student’s t-tests for two-group comparisons, and one-way or two-way analysis of variance (ANOVA) tests were used for normally distributed data, followed by post hoc analyses for multiple group comparisons, as noted in the legends. p < 0.05 was considered statistically significant.

## Data Availability

All data that support the findings of this study are available from the corresponding authors upon reasonable request.

## Code Availability

All computer code generated during and/or used in the current study is available from the corresponding authors on reasonable request.

## Acknowledgments

The authors thank Lindsey Butler for mouse colony management. This work made use of the NUFAB facility of Northwestern University’s NUANCE Center, which has received support from the SHyNE Resource (NSF ECCS-2025633), the IIN, and Northwestern’s MRSEC program (NSF DMR-2308691). MicroCT imaging work was performed at the Northwestern University Center for Advanced Molecular Imaging (RRID:SCR_021192) generously supported by NCI CCSG P30 CA060553 awarded to the Robert H Lurie Comprehensive Cancer Center. Some schematic illustrations used materials created by BioRender.com. This work was funded by the Querrey-Simpson Institute for Bioelectronics (MW, YY, AIE, AVG, YW, JG, LZ, JL, MK, JK, YH, JAR); NINDS/BRAIN Initiative 1U01NS131406, NIMH R01MH117111, NINDS R01NS107539, 2021 One Mind Nick LeDeit Rising Star Research Award (YK); Shaw Family Pioneer Award, Center for Reproductive Science, Feinberg School of Medicine (JMC); NIMH R00MH120047, Simons Foundation grant 872599SPI, Alfred P. Sloan Foundation grant SP-2022- 19027 (LP); NC State University Start-up fund 201473-02139 (AVG); and the Christina Enroth-Cugell and David Cugell fellowship (MW).

## Author contributions

Conceptualization: MW, YY, YK, JAR; Methodology: MW, YY, AIE, JZ, AVG, XL, YK, JAR; Theoretical simulations: XL, KZ, MW, WZ, AVG, YH; Investigation: MW, YY, JZ, AIE, AVG, XL, KZ, YW, JG, LZ, JL, MR, HY, MK, HZ, ML, JK, KT, SC, AB, CHG, JMC, LP, YH, YK, JAR; Software: AVG, HY, YW, MR; Formal analysis: MW, JZ, XL, KZ, HY; Validation: MW, JZ, AIE; Data curation: JZ, LZ, HY, AIE; Visualization: JZ, MW, YY, AIE, XL, KZ, YW, JG, JL, MR; Supervision: YY, YH, YK, JAR; Funding acquisition: YH, YK, JAR; Writing-original draft: MW, YY, JZ, AIE, AVG, XL, KZ, YK, JAR; Writing-review & editing: MW, YY, JZ, AIE, AVG, XL, KZ, JL, JMC, LP, YH, YK, JAR. MW, YY, JZ, AIE, AVG, KZ, XL contributed equally to this work.

---

**Extended Data Fig. 1.**
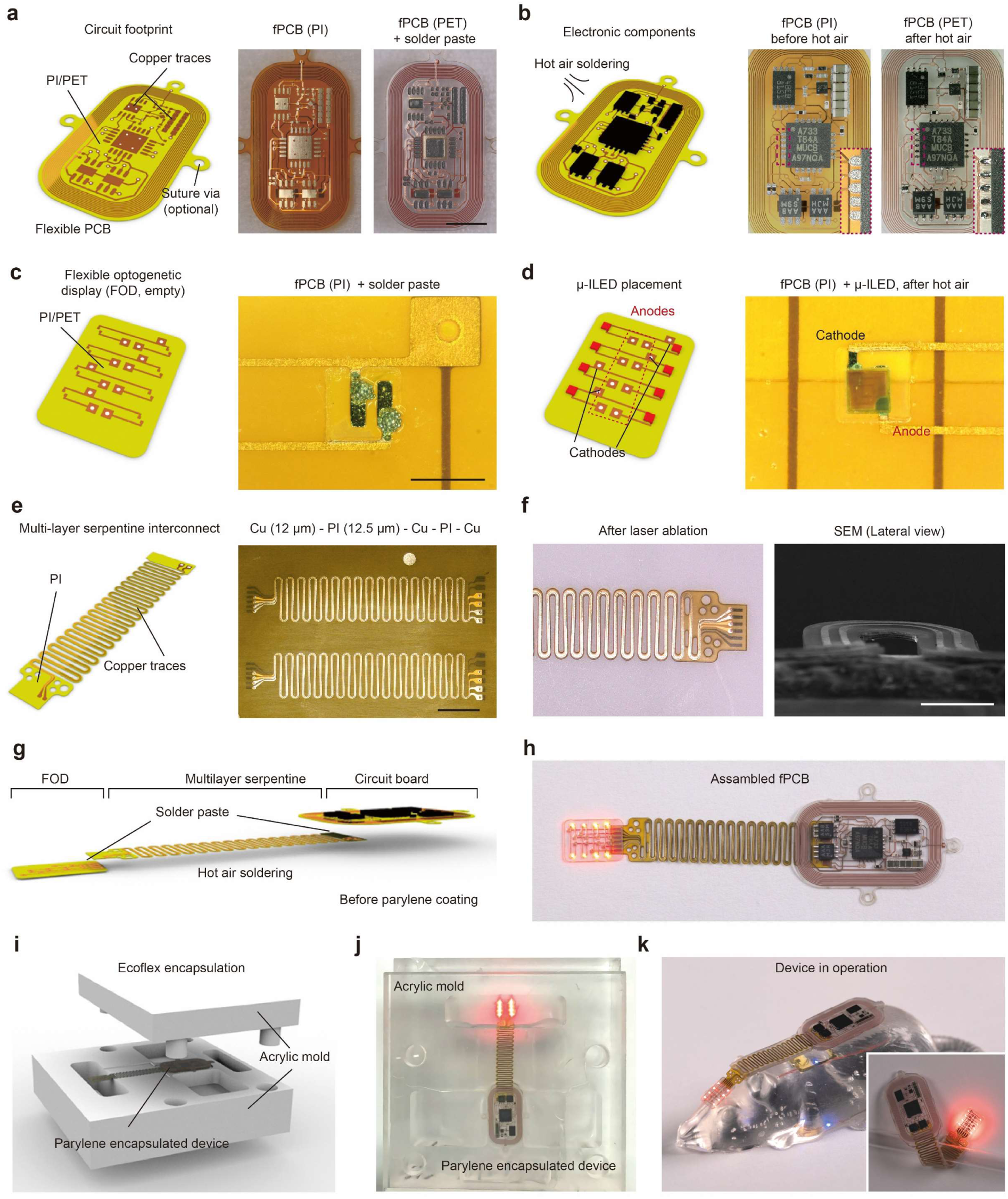
Device fabrication and assembly. (a) Left, schematic illustration of the circuit base of the electronic module of the device using three-layer flexible printed circuit board (fPCB). Right, images of the fPCB for the electronic module. Scale bar: 5 mm. (b) Left, schematic illustration of the assembly of electronic components on the fPCB of the electronic module using hot-air soldering. Right, images of the electronic module after hot-air soldering of electronic components. (c) Left, schematic illustration of the circuit base of the FOD using three-layer fPCB. Right, magnified view of the soldering site for the μ-ILED with solder paste. Scale bar: 500 µm. (d) Left, schematic illustration of the FOD after hot-air soldering of the μ-ILEDs. Right, magnified view of the soldered μ-ILED. (e) Left, schematic illustration of the serpentine traces of the device using a five-layer fPCB. Right, image of the serpentine traces on the fPCB. Scale bar: 5 mm. (f) Left, image of the serpentine traces after laser ablation from the fPCB substrate. Right, scanning electron microscopic (SEM) image of the serpentine traces from lateral view. Scale bar: 300 µm. (g) Schematic illustration of the final assembly of the electronic module, serpentine traces, and FOD. (h) Image of the final device after assembly. (i) Schematic illustration of the mold used for final silicone (Ecoflex 00-30) coating after parylene-C encapsulation. (j) Photograph of the device during mold casting of silicone. (k) Image of a fully encapsulated device ready for in vivo experiments.

**Extended Data Fig. 2.**
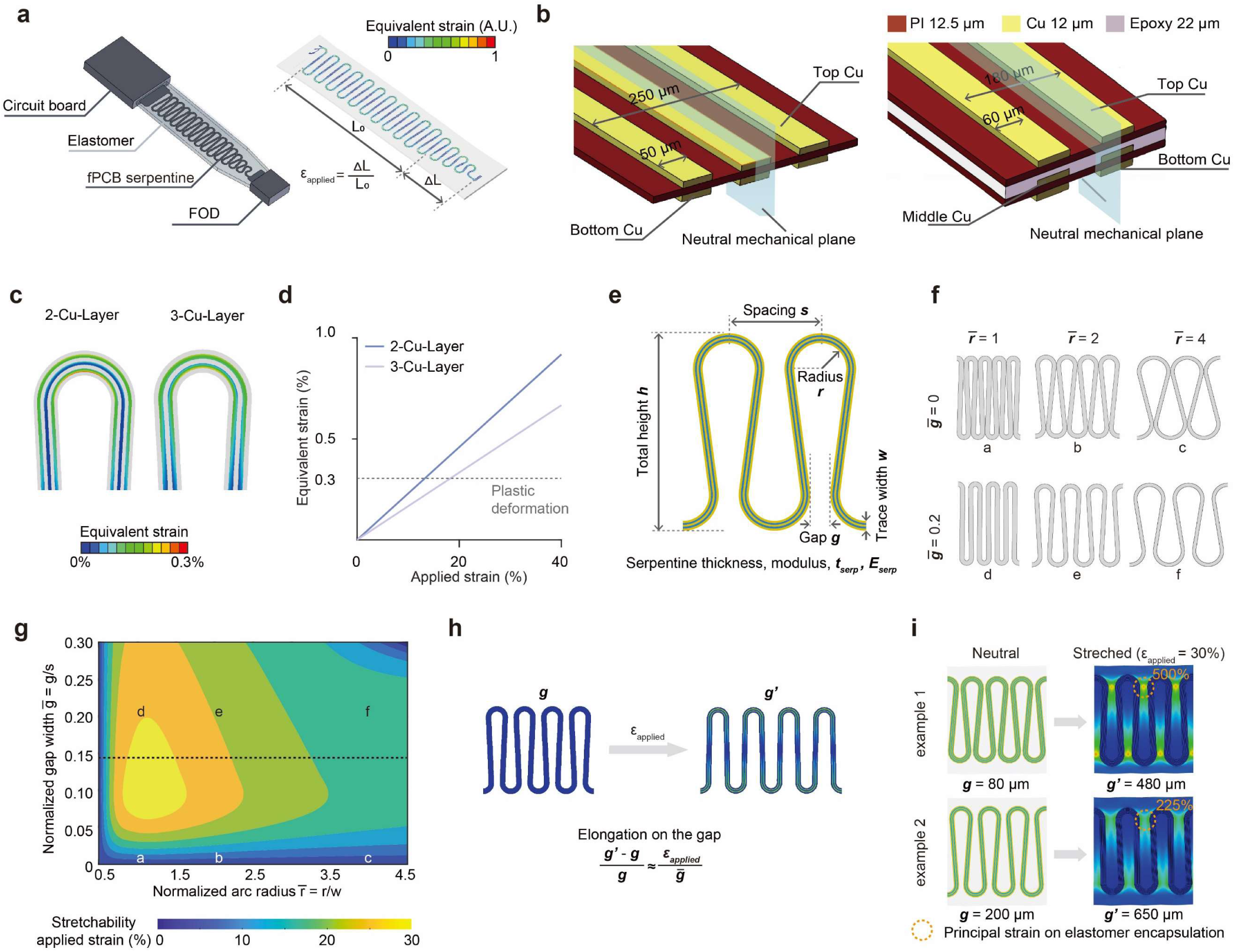
Geometric optimization of serpentine interconnect. (a) Left, geometric model of structures surrounding the serpentine interconnect. Right, calculation of the applied strain on the serpentine and equivalent strain on the Cu-based serpentine conductive traces. (b) Layered structure of the serpentine materials relative to the neutral mechanical plane. Left: 2-Cu-layer design; right: 3-Cu-layer design. (c) Equivalent strain distribution on copper traces with 13% applied strain. (d) When a certain strain is applied to the serpentine interconnect, a certain copper unit on the trace shows the highest equivalent strain among all units. Summary graph showing this highest equivalent strain versus applied strains. Equivalent strain for copper plastic deformation equals 0.3%. (e) Geometric parameters that affect the stretchability of the serpentine interconnects. (f) Serpentine design examples with different gap widths and radius. (g) Contour map summarizing the stretchability as a function of normalized gap width and radius. Stretchability: maximal applied strain without causing plastic deformation on copper-based conductive traces. (h) Average elongation of the gap when the serpentine interconnect is stretched. (i) Elongation of the gap results in principal strain on the elastomer, causing potential encapsulation defects.

**Extended Data Fig. 3.**
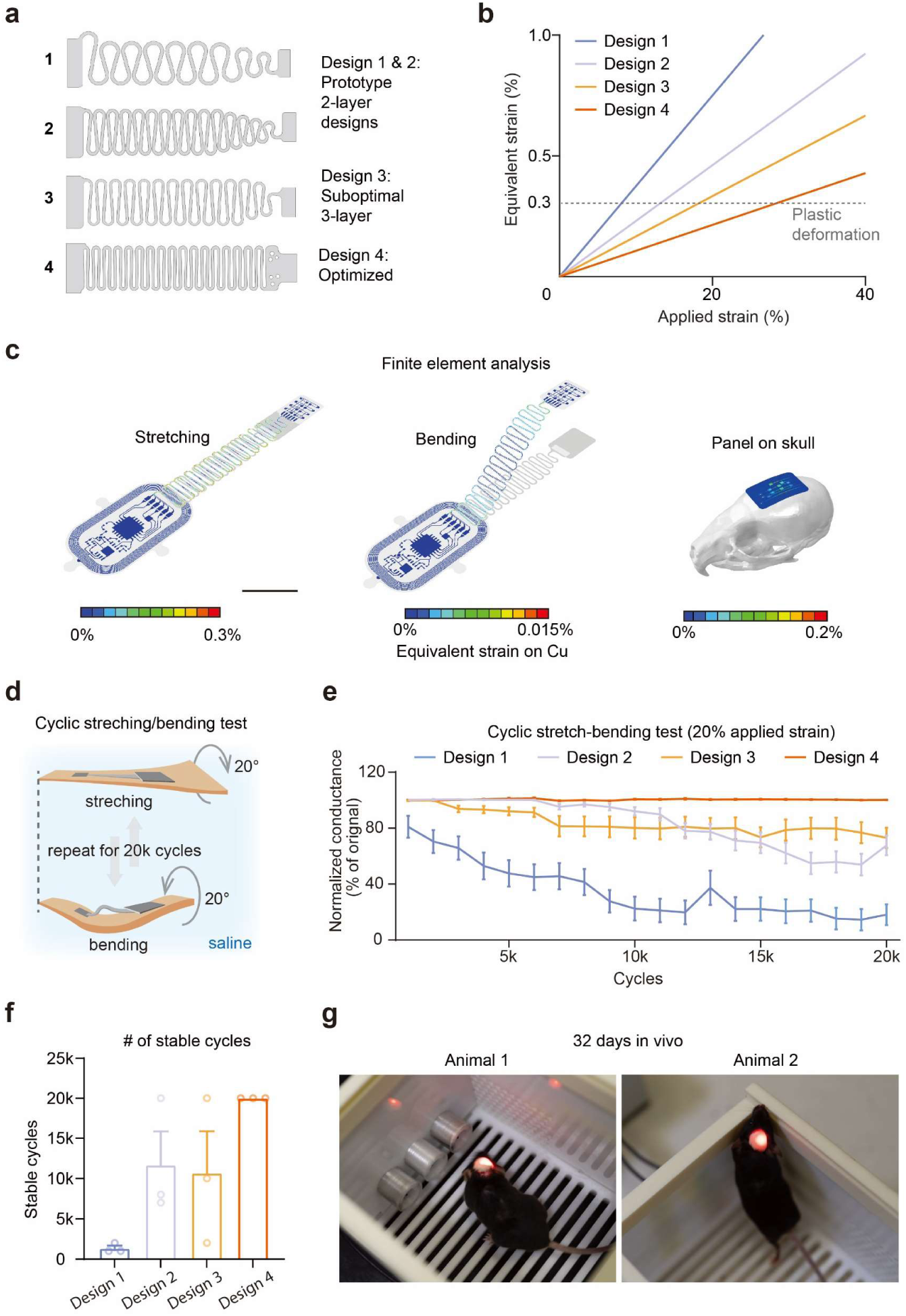
Numerical modeling results and experimental characterizations for four serpentine designs. (a) Design 1 & 2: prototypes, not guided by numerical modeling; Design 3, optimized layer structure with preliminary modifications of gap width and connecting regions; Design 4, optimized layer structure and geometry guided by numerical modeling. (b) Equivalent strain on the Cu-based conductive traces versus applied strain on the serpentine interconnect for the four designs. Dashed line: Equivalent strain threshold for plastic deformation on the copper traces. (c) Finite element analysis (FEA) of equivalent strain on the optimized serpentine interconnect under stretching (left) and bending (middle). Right, the equivalent strain on the FOD when flexed to the curvature of the skull. Scale bar: 10 mm. (d) Schematic illustration of the benchtop validation of the modeling outcomes using cyclic stretching and bending tests of devices integrated on artificial skin immersed in saline. (e) Summary data showing normalized conductance loads on individual μ-ILEDs for 20k cycles in cyclic stretching and bending tests. n = 16 - 24 μ-ILEDs from 3 devices for each design. (f) Summary data for stable cycles where the devices maintain constant resistance during stretching and bending. n = 3 devices. (g) Images showing devices in operation in the Bpod chamber, 32 days after implantation.

**Extended Data Fig. 4.**
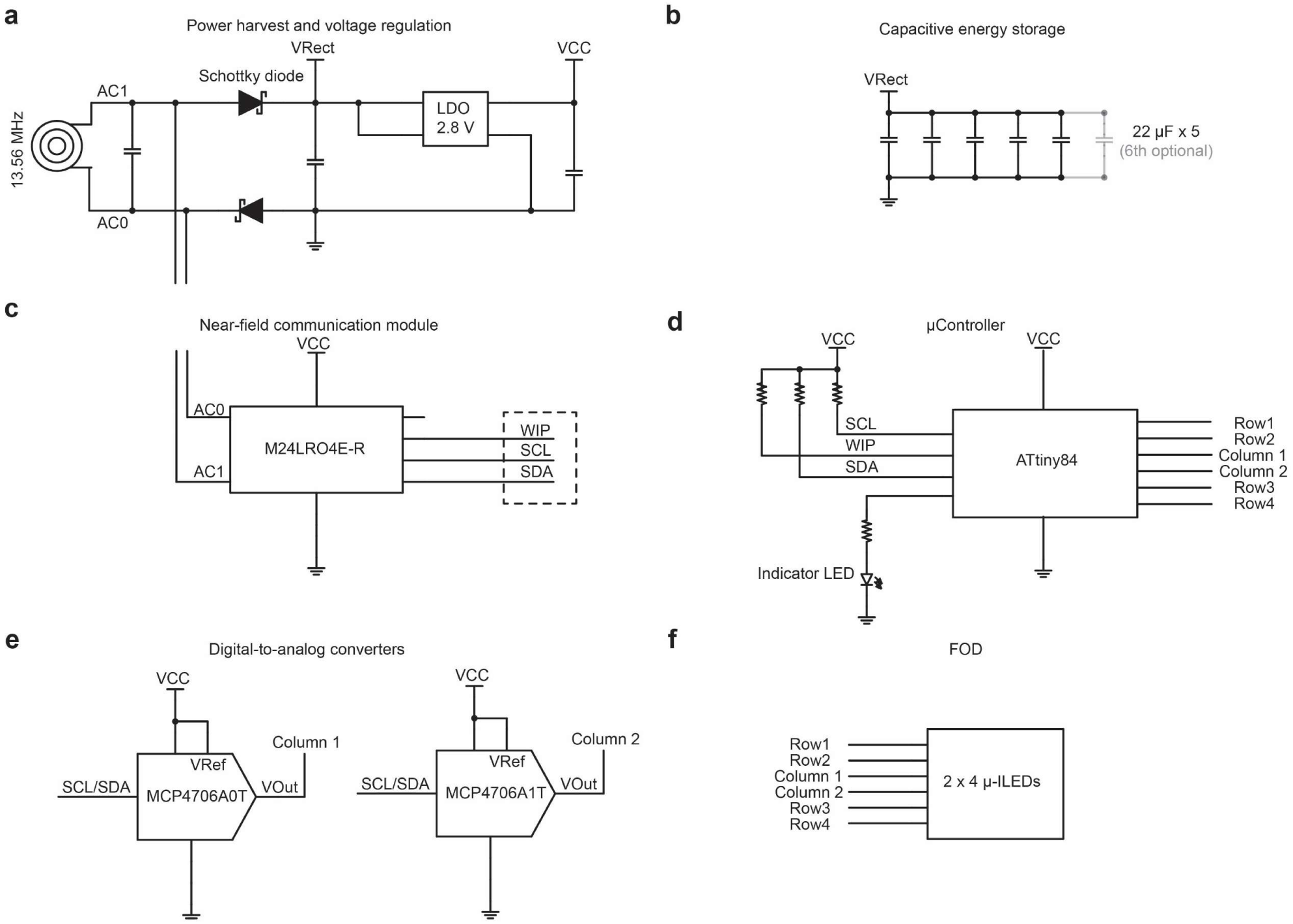
Electronic modules. (a) Schematic illustrations showing electronic circuits for wireless power harvesting and voltage regulation. (b) Capacitor bank for energy storage and discharge. (c) Near-field communication module for real-time programming. (d) Micro-controller circuit for parameter (order, frequency, duty cycle) control. (e) Digital-to-analog converters for intensity modulation. (f) FOD. Essential components and input/output pins are labeled on the schematics.

**Extended Data Fig. 5.**
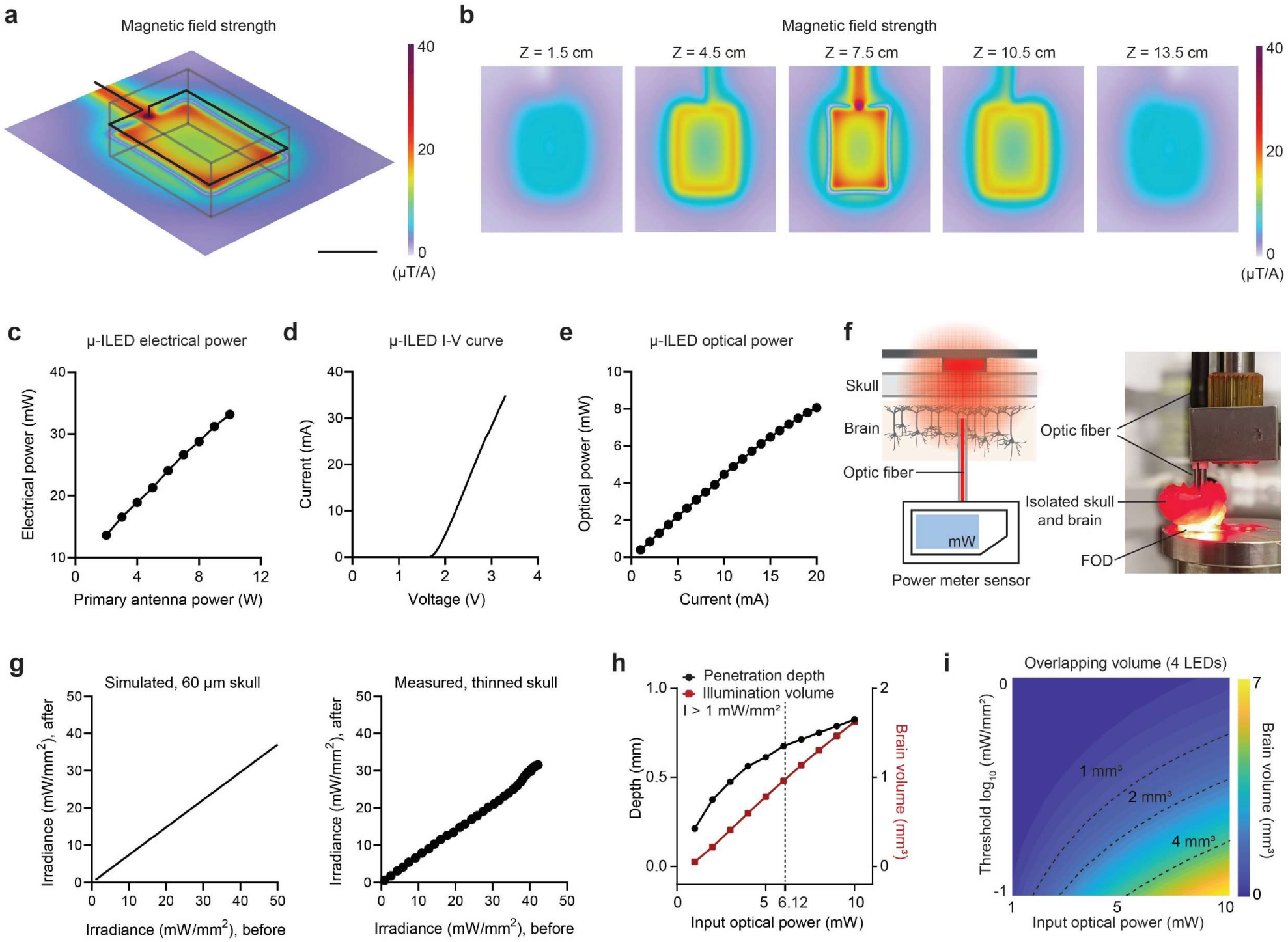
Characterizations and validations for irradiance distribution profiles. (a) Simulated magnetic-field-intensity distribution at the central plane of a behavioral cage (dimensions, 20 cm (length) × 14 cm (width)) with a double-loop antenna at heights of 3 cm and 6 cm. Scale bar: 10 cm. (b) Simulated magnetic-field-intensity distribution at different heights in the Bpod behavioral cage. (c) The total electrical power of the μ-ILED with respect to the primary antenna power. (d) The I-V characterization for the μ-ILED. (e) The optical power of the μ-ILED with respect to the input current. (f) Schematic illustration (left) and image (right) of the experimental setup for measuring light attenuation through the skull and brain tissue. (g) Simulated (left) and measured (right) results of light attenuation through the skull. A 60 µm layer of skull was used in the numerical model. A piece of thinned skull was used for measurement. (h) Illumination volume and penetration depth as a function of the input irradiance of the red μ-ILED (628 nm). Threshold intensity, 1 mW/mm^2^. (i) Contour map plot showing total overlap volume from 4 μ-ILED co-activation for different input optical power and intensity thresholds for opsin variants.

**Extended Data Fig. 6.**
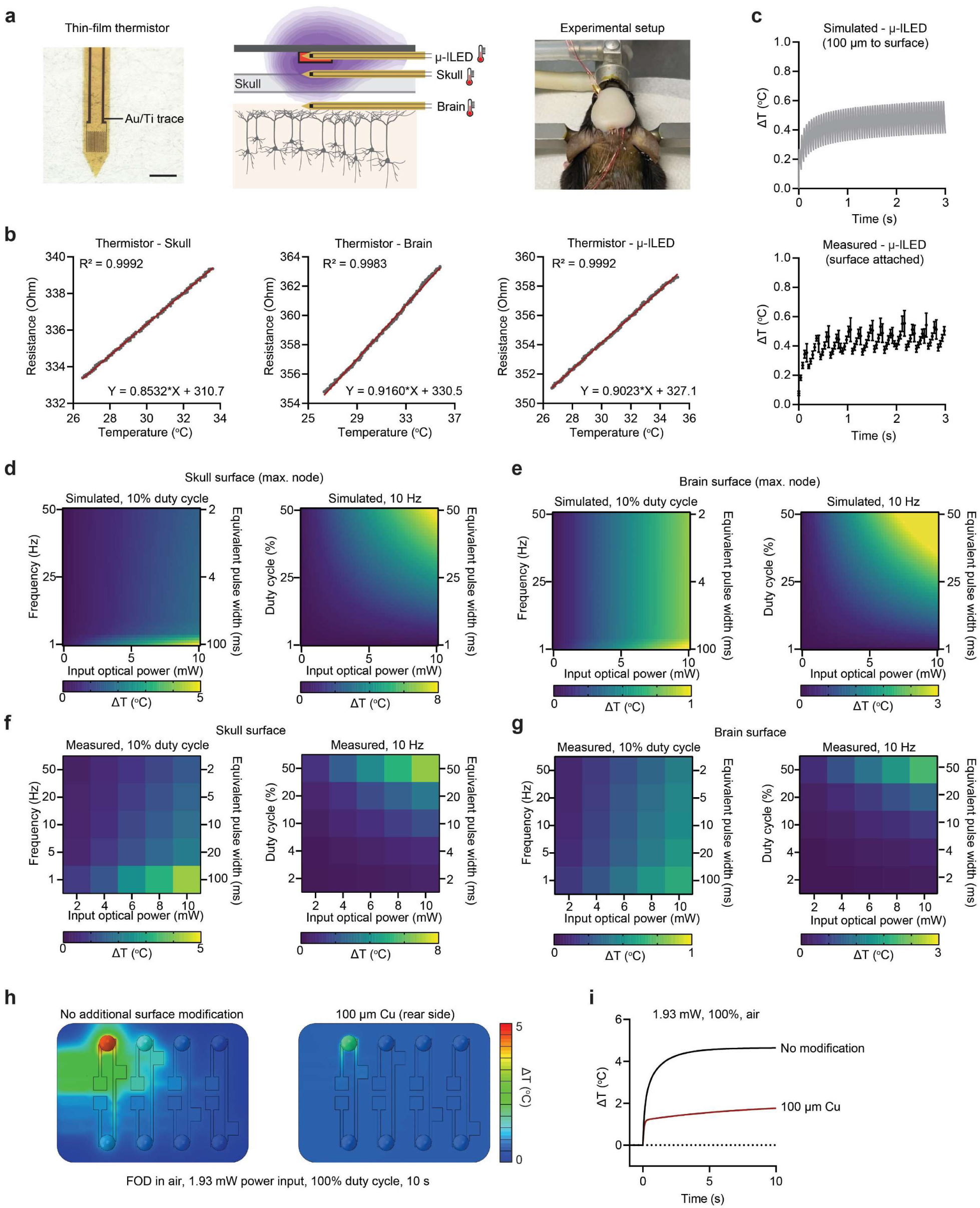
Numerical and experimental assessments of heat accumulation to guide stimulation protocols. (a) Left, image of micro-fabricated thermistor for measuring temperature; middle, schematic illustration of the experimental setup for measuring heat accumulation in the structures surrounding the optical- neural interface; right, image of experimental setup, with wires connected to the thermistors to collect resistance value. Scale bar: 500 µm. (b) Calibration curves of thermistors used in this study for measuring the temperature increase on the surface of skull (left), on the surface of brain (middle), and below the µ-ILED (right). (c) Simulated (Top) and measured (bottom) results of temperature increase below the µ-ILED for 3 s μ-ILED operation. The temperature increases in a 20 × 300 × 300 µm^3^ volume, 100 µm below the μ-ILED surface, was output from the numerical model. The thermistor was manually placed and adhered to the μ-ILED surface, followed by a complete procedure of μ-ILED array fabrication. (d) Left, simulated maximal temperature increases on the skull surface during μ-ILED operation with 10% duty cycle at varying frequencies. Right, same as left, but for varying duty cycles at 10 Hz. The finite element node with maximal temperature increase (max. node) was selected for plotting. (e) Same as (d), but for brain surface. (f) Left, measured temperature increases on the skull surface during μ-ILED operation with 10% duty cycle at varying frequencies. Right, same as left, but for varying duty cycles at 10 Hz. (g) Same as (f), but for brain surface. (h) Heatmap showing simulated heat production during a single μ-ILED operation in air at 100% duty cycle and 1.93 mW power for 10 s. Left, fPCB array patch without modification. Right, fPCB array patch coated with 100 µm Cu on the rear side. (i) Simulated temperature increases with or without device surface modification with 100 µm Cu.

**Extended Data Fig. 7.**
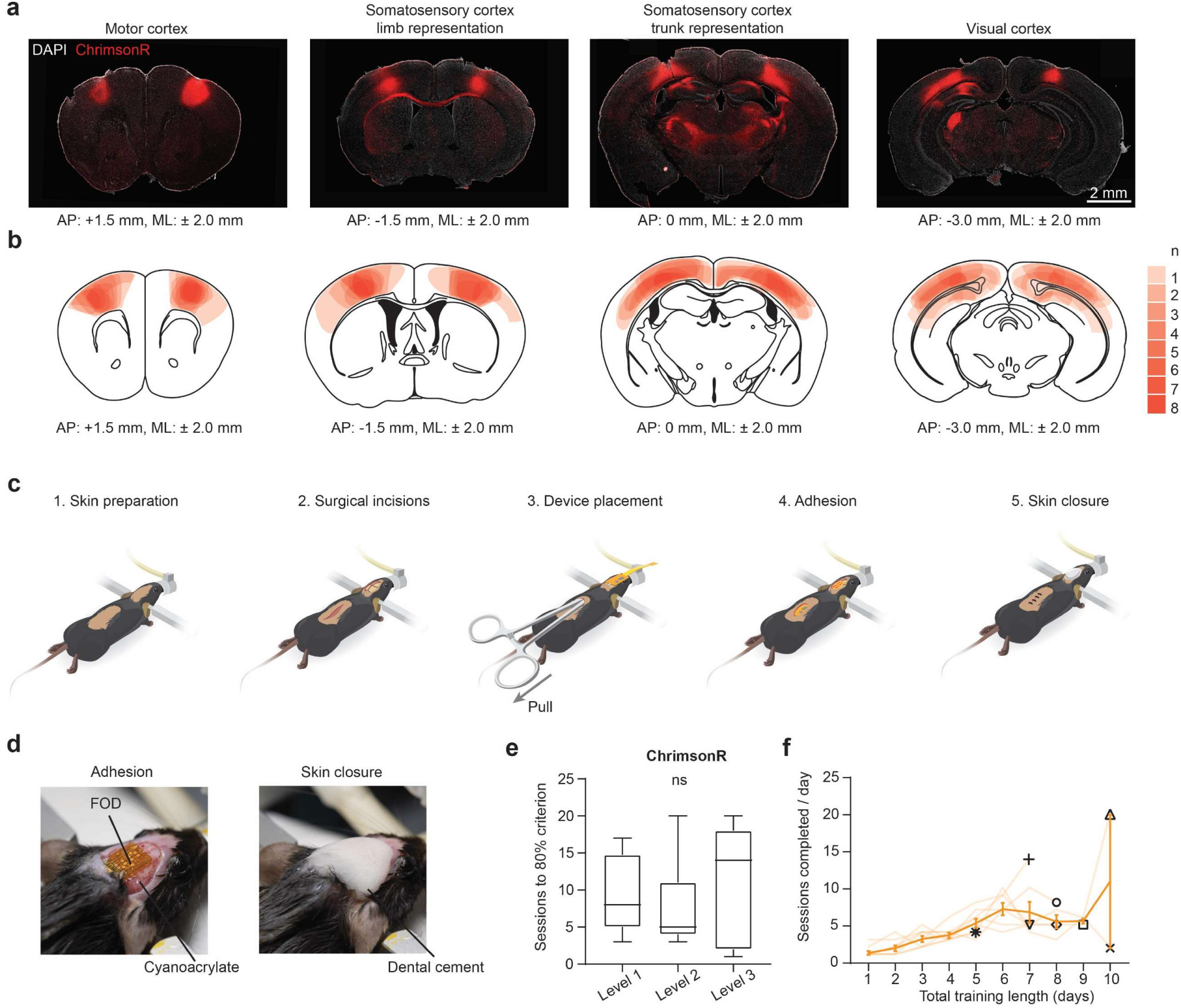
Opsin expression, surgical procedures, and behavioral performance after implantation. (a) Representative images of ChrimsonR expression in the targeted cortical regions from one mouse. (b) Heatmap showing the spread of viral expression across cortical regions. n = 8 animals. (c) Schematic illustration of critical steps of surgical implantation. Detailed step-by-step surgical procedures are documented in our published protocols^42^. (d) Photographs showing the FOD attached to the mouse’s skull with cyanoacrylate adhesives and fixation with dental cement. (e) Number of sessions of 100 trials each across three levels of the task to reach criterion of 80% success rate for animals expressing ChrimsonR (session median Level 1-8, Level 2-5, Level 3-14). p = 0.7061, F (2, 19) = 0.3545; Level 1, n = 8 animals; Level 2, n = 7 animals; Level 3, n = 7 animals. (f) Summary data showing the total training length to complete Levels 1-3 and the number of sessions per day. Dark orange, group average; pale orange, individual trajectories; symbols mark the day individual animals reached criterion.

**Extended Data Fig. 8.**
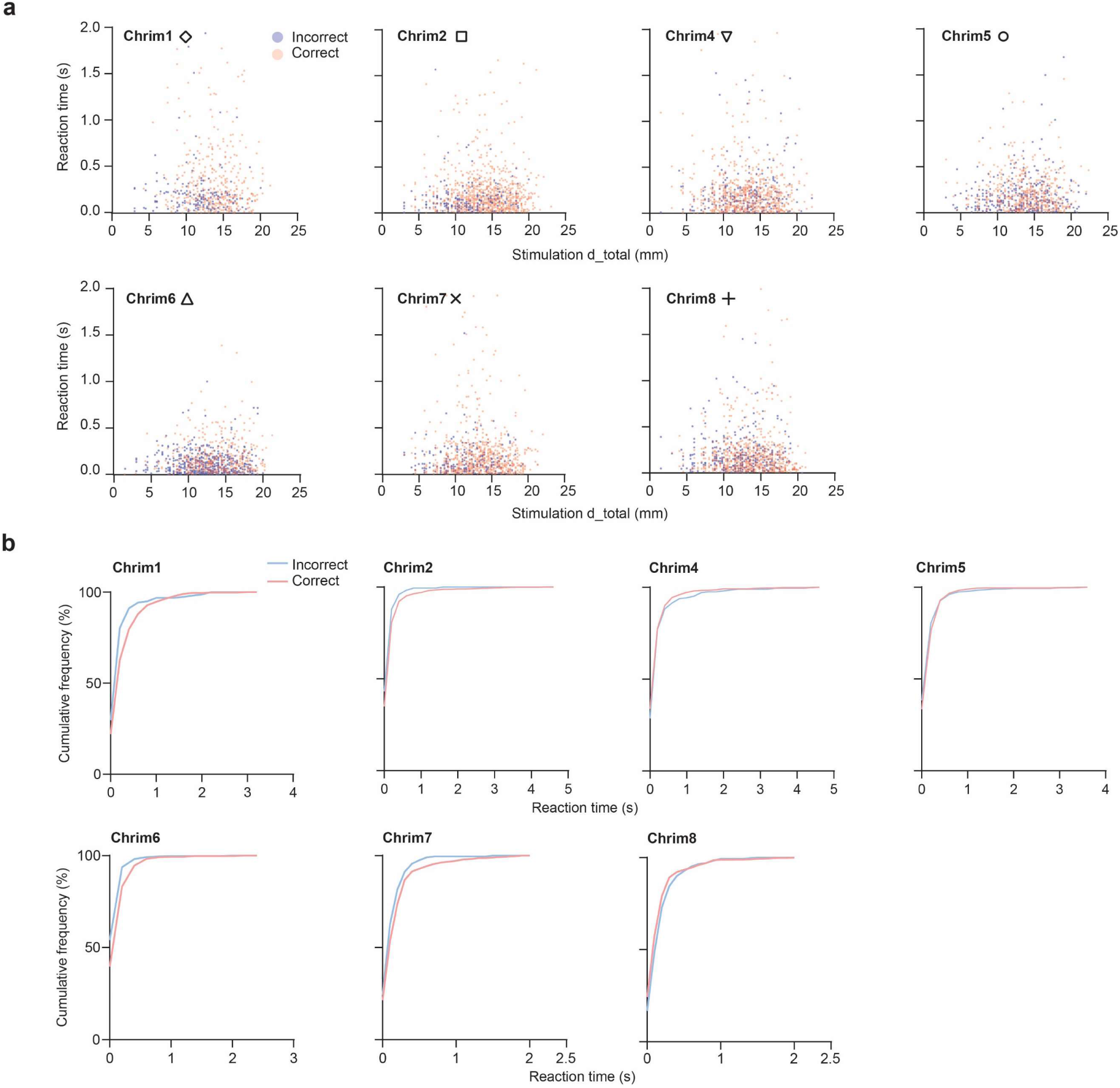
Reaction time analysis during operant learning behaviors. (a) Scatter plots showing reaction times in all trials for Level 3 task for individual animals as a function of total cortical distance between stimulated digits. (b) Same as (a), but shown as cumulative distributions of reaction times.

**Extended Data Fig. 9.**
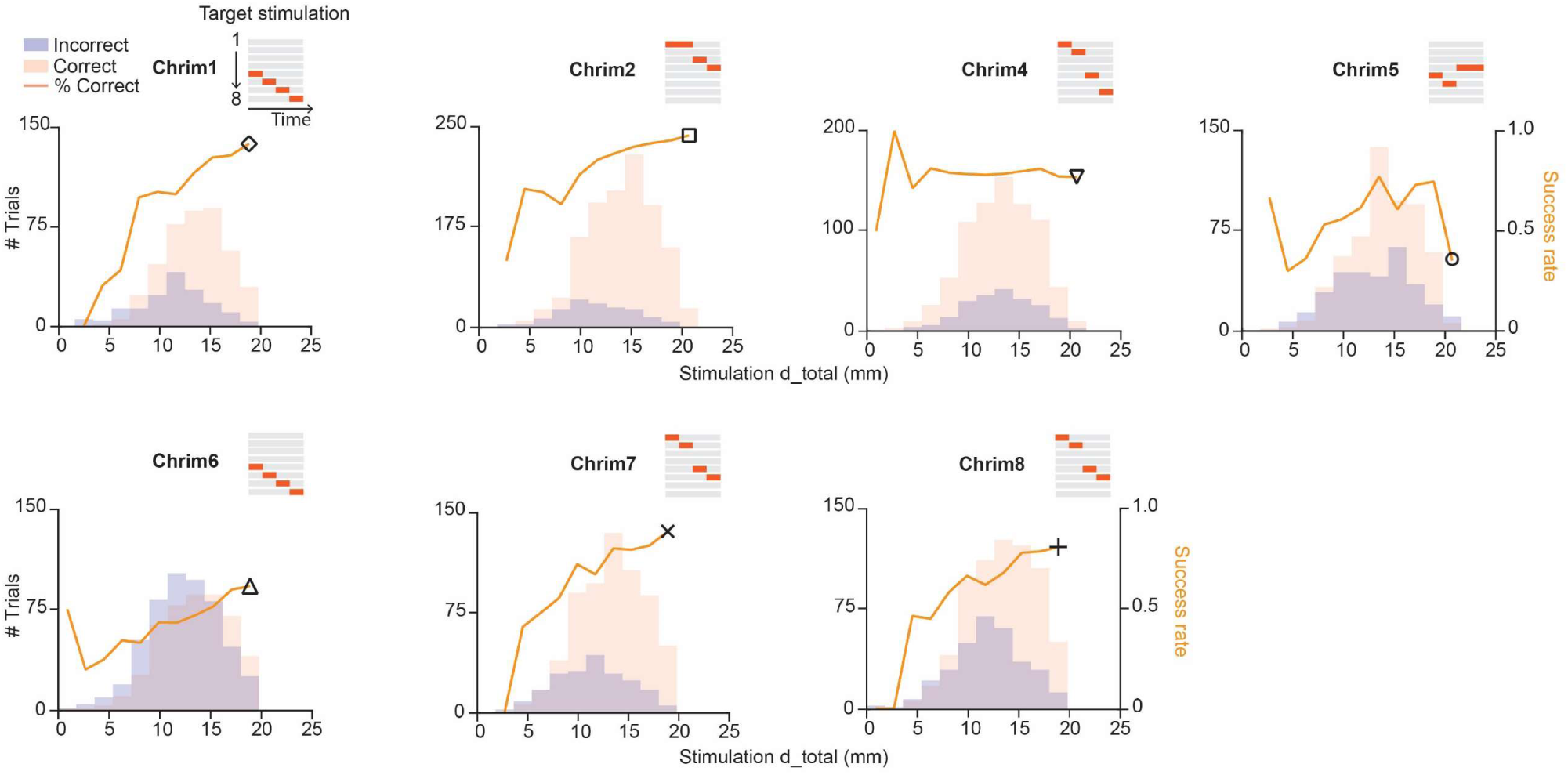
Spatial distance analysis of behavioral trajectories for individual animals. Line plots showing success rate as a function of spatial distance for 7 individual animals in the Level 3 task. The number of trials with correct or incorrect choices in each bin of spatial distance is plotted in the histogram.

**Extended Data Fig. 10.**
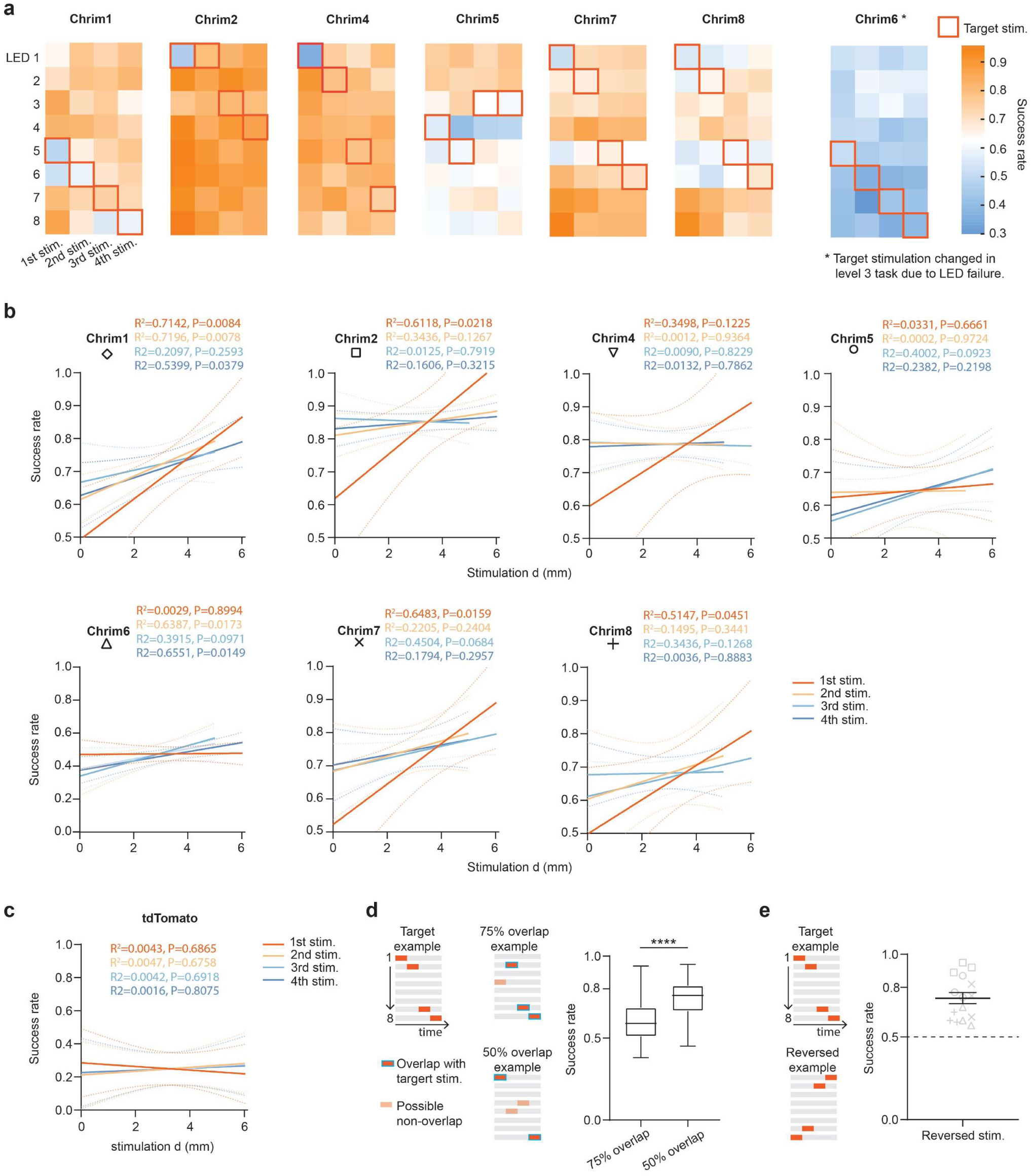
Digit-based analysis of behavioral trajectories and probing experiments with similar or reversed sequences. (a) Left, heatmap showing success rate for randomized non-target sequences grouped by specific stimulation locations at the first to the fourth stimulation digit in seven ChrimsonR expressing animals. Red squares indicate the target stimulation sequence for each animal. Right, same but for one ChrimsonR-expressing animal that experienced µ-ILED failures during training, requiring target sequence reassignment (serving as an internal control). (b) Pearson’s correlation analysis of spatial distance and success rate based on stimulation digit for individual animals. (c) Pearson’s correlation analysis of averaged spatial distance and averaged success rate based on stimulation digit for all animals expressing tdT, showing no relationship between distance and success rate for any digit. (d) Left, schematic illustration of probing sequences with 75% (3 stimulation digits) and 50% similarity (2 stimulation digits) to the target sequences. Right, summary data showing the median and quartile success rate of all probing sessions from all animals. Two-sided unpaired t-test, p < 0.0001. 75% overlap: n = 75 sessions from 5 animals; 50% overlap: n = 41 sessions from 5 animals. (e) Left, schematic illustration of probing experiments with reversed sequences. Right, summary data showing success rate of all probing sessions from all animals. n = 15 sessions from 5 animals.

